# A quantitative narrative on movement, disease and patch exploitation in nesting agent groups

**DOI:** 10.1101/791400

**Authors:** Wayne M. Getz, Richard Salter, Krti Tallam

## Abstract

Animal relocation data has recently become considerably more ubiquitous, finely structured (collection frequencies measured in minutes) and co-variate rich (physiology of individuals, environmental and landscape information, and accelerometer data). To better understand the impacts of ecological interactions, individual movement and disease on global change ecology, including wildlife management and conservation, it is important to have simulators that will provide demographic, movement, and epidemiology null models against which to compare patterns observed in empirical systems. Such models may then be used to develop quantitative narratives that enhance our intuition and understanding of the relationship between population structure and generative processes: in essence, along with empirical and experimental narratives, quantitative narratives are used to advance ecological epistemology. Here we describe a simulator that accounts for the influence of consumer-resource interactions, existence of social groups anchored around a central location, territoriality, group-switching behavior, and disease dynamics on population size. We use this simulator to develop new and reinforce existing quantitative narratives and point out areas for future study.

**Author summary:** The health and viability of species are of considerable concern to all nature lovers. Population models are central to our efforts to assess the numerical and ecological status of species and threats posed by climate change. Models, however, are crude caricatures of complex ecological systems. So how do we construct reliable assessment models able to capture processes essential to predicating the impacts of global change on population viability without getting tied up in their vast complexities? We broach this question and demonstrate how models focusing at the level of the individual (i.e., agent-based models) are tools for developing robust, narratives to augment narratives arising purely from empirical data sources and experimental outcomes. We do this in the context of nesting social groups, foraging for food, while exhibiting territoriality and group-switching behavior; and, we evaluate the impact of disease on the viability of such populations.

## 1 Introduction

Disease ecology is a rich field of study that synthesizes *inter alia* demographic, epidemic, behavioural, movement, spatial and community processes in an ecological setting [1–6]. Thus, understanding the impact of particular diseases on wild populations in ecological settings is a challenge that requires a comparison of the spatial and community structure of these focal populations, as they might arise in the absence versus the presence of the causative agents of the diseases of interest. One can undertake such studies in the laboratory at the microcosm scale [7], but for populations of most vertebrates such microcosm studies are not applicable, and we need to resort to the mesocosm level [8] or use alternative methods to obtain insights. One of these methods is to build simulation models that incorporate demographic, spatial, movement, and other relevant behavioral and community processes, and then compare simulations of these models across various scenarios [9] or outcomes with and without epidemiological processes added to the mix. We take this latter approach and our primary aim is to develop a tool that can be used to study the impact of disease on colonial populations and metapopulations [10]. This tool must be comprehensive enough to include critical processes needed to address questions that will extend our current knowledge in significant ways. Its application, of course, is limited by the structure of the model. Thus, in building the model, we are guided by principles related to concepts of “appropriate complexity modeling” [11, 12].

The standard approach to modeling disease outbreaks caused by directly-transmitted pathogens is through systems of differential or difference equations that model the number or density of susceptible (S), infectious (I), and recovered or removed (R) individuals [13, 14]. These formulations may either be deterministic or stochastic, and elaborated to include time-delays [15], demographic (births, deaths, and recruitment [16]), metapopulation [17] and other spatial processes [18]. Examples of these elaborations can be found in the context of modeling, *inter alia*, influenza [19, 20], measles [21], SARS [22, 23], and Ebola [24–28] outbreaks.

Systems approaches to modeling disease limit our ability to consider individual-level traits, unless these traits can be reasonably well expressed by dividing the population into a limited number of subgroups, such as groups of individuals infected with different strains of pathogens [29]. Including individual-level variation in disease models, however, requires an agent-based approach [30], which then allows both individual behavior (e.g., relevant to evaluating the probability of pathogen transmission) and pathogen exposure history (e.g., relevant to evaluating the immunological state and hence the susceptibility of an individual) to be incorporated into the model. It also allows one to trace the number of individuals the each pathogen infects [25] and, hence, to identify individuals that may be acting as super-spreaders [31].

Here, we develop an agent-based formulation that has application to studying disease outbreaks in wildlife populations whose state of health—and hence, susceptibility—is impacted by various factors, including social factors related to group living and the structure of social networks [32], ecological factors related to foraging success, and the movement behavior of individuals (which influences exposure to pathogens). The model also includes a periodic return to a colony or nest cell, which implies that the model has both a population-level and a group-association level (core or household) structure. Such models have applications in animal production systems (e.g., managing foot and mouth disease [33]), wildlife conservation [2], and disease management at the human-animal interface [34]. The latter involves diseases such as anthrax [35], bovine tuberculosis [36], and brucellosis in cattle and wild ungulates [37], chronic wasting disease in deer [38], and toxoplasmosis in rats, cats and humans [39].

Rather then following a formulaic ODD (Overview, Design concepts, and Design details; [30, 40]) or extended ODD [41] prescription in presenting our model, we take a more expository approach by first crafting a broad outline of our model’s general features. In addition, to keep the paper reader-friendly, we relegate the mathematical details of our model to a Technical Methods section at the end of the paper and the algorithmic details to a supporting online file (SOF). In the Results section, we report the results of simulations of several illustrative scenarios, and in the Discussion section we present the findings in the form of a quantitative narrative that provides insights into the role different processes play in creating the population patterns observed in our simulations.

At this point, a few brief remarks on quantitative narratives in disease ecology are in order. Unlike the canonical theories of physics—with kinematics, electromagnetism, general relativity, and quantum mechanics providing prime examples, ecology has no fundamental theories and associated mathematical formulations that can be used to develop an ecological epistemology. Rather, the complexity of the field induced by the plethora of processes involved requires that we focus only on a limited number of these processes when building models, where the selection of process depends on the questions at hand [11]. Our understanding of these processes requires an analysis of associated quantitative formulations, but the soundness of the arguments cannot be adjudicated by comparing, say, the accuracy of Mercury’s orbit as predicted by Newtonian versus Einsteinian physics. Rather, ecological epistemology is woven from a collection of quantitative narratives that provide current best explanations for, or fits to, observed empirical and experimental data. While statistical null models are use to develop consilient empirical and experimental narratives, dynamic null models are used to develop quantitative narratives that are consilient with empirical and experimental narratives (Fig. 1). Such dynamic null models have a Goldilocks quality in the sense of containing just the right amount of biological process detail [42]: too little and model simulations are rejected as providing adequate descriptions of observed data; too much process detail and the null-model-parsimony principle, as discussed by Gotteli and Graves ([42], p. 6), is violated.

**Fig 1.**
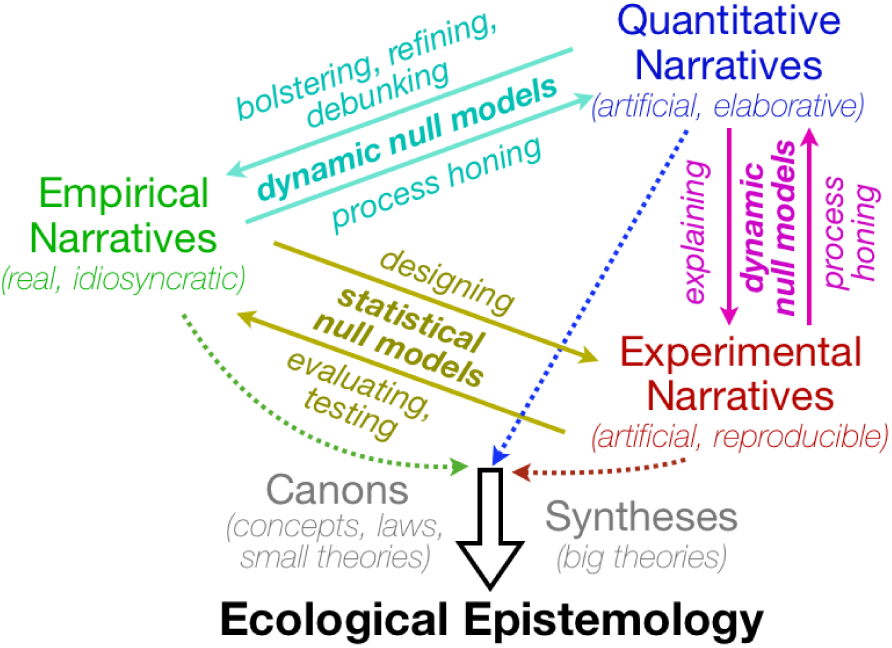
A sketch of the role of empirical, experimental and quantitative narratives in the formulation of canons and construction of syntheses in the development of an ecological epistemology.

In ecology, quantitative narratives, derived from the Gause-Lotka-Volterra competition model, were used to construct the concept of a niche [43]. This narrative begins with insights into competitive exclusion [44, 45], supported by additional quantitative ideas relating to niche breadth and overlap [46, 47]. In epidemiology, quantitative narratives, derived from analyses of SEIR epidemic models (S: susceptible; E: exposed but not yet infectious; I: infections; R: removed—that is, recovered with immunity or dead) [13], have been used to identify critical differences in the emergence of outbreak thresholds depending on whether pathogen transmission is dependent on the density or frequency of infected individuals in the population [48, 49]. Other quantitative narratives in epidemiology that incorporate aspects of the movement of individuals [50] or particular ecological processes have been used to develop canonical theories [51–55]; however, these are limited in scope and generality by the analytical difficulties of studying dynamical systems, as well as networks that include more than one ecological or behavioral process. The challenge here is to keep the models somewhat general and broadly applicable, while including several key processes such as: spatial demography; consumer-resource ecology with associated reproduction, growth, stress, and mortality processes; movement and contact behavior; and SEIR epidemiology, in ways that are simple enough to interpret the effects of each process. Sensitivity analyses provide one type of approach to separating out effects [56–59]. Another is to use methods relating to the practice of appropriate complexity modeling [11, 12].

## 2 Methods

The model we present here only incorporates the focal processes of interest and those needed to attain a level of realism required to adequately address the questions at hand. Thus, for example, we do not include agent age structure in the model even though age structure is known to have important effects on population growth patterns [60], persistence [61], and epidemics [62]. Agent age structure would certainly add complexity to our narratives; but, whether its impacts are critical (i.e., it qualitatively changes the structures we explain) or only elaborative (i.e., has quantitative but not qualitative implications regarding the questions asked) is a issue to be examined in future studies. For clarity, we enumerate below—without mathematical details—the processes incorporated in our model, leaving such details to the Technical Methods Section at the end of the paper. Aspects of the Numerus Model Builder construction of the model and the generated NovaScript (extended JavaScript language) implementation are provided in a Supporting Information file.

### 2.1 Model Outline

The model is formulated as a set of agents (individuals) moving over a cellular array. The agents themselves can be differentiated with respect to type (e.g., sex), can undergo discrete state transitions (e.g., disease state) and can also change continuously over time with respect an agent-level state vector. In our case, the latter contains the variables “mass” (also a surrogate for fitness) and “stress” (which accumulates when the individual is unable to meet its resource needs each time period). The cells in the cellular array may also have relatively complex states that, in our model, include a list of the agents in each cell and the level of a dynamically responding resource, which determines the environmental context for the agents. Movement of agents occurs under the assumption that, apart from reproduction, ecological and epidemiological processes are independent of agent type. More specifically, in our model, the following objects and concepts apply:

#### Simulation world

This is the set of agents moving over a hexagonal cellular array (we use a toroidal topology to eliminate boundary effects).

#### Cells

These are spatial extents (or patches) that contain resources (e.g., plants, fruits, seeds), host at most one colony site (i.e., are a nest or colony cell), and can be occupied by one or more agents who either pass through in a single time period or remain for some time during which they exploit the dynamically changing (through both growth and extraction) resources.

#### Time

The action takes place over discrete time steps during which individuals (agents) move through one or more cells, stay in a cell to exploit resources, or return to their colony cell at regular intervals of time where they may reproduce.

#### Agents

Each agent has a biomass (which is a surrogate measure of its fitness), a stress level, and a sex determined at birth. Females may reproduce on regular returns back to their colonies by, essentially, fissioning into two (the parent retains the majority of the biomass, while the offspring acquires the minority of biomass, with some loss due to the cost of reproduction).

#### Agent association groups (colonies)

Each agent belongs to one of several association groups called colonies. These colonies have a home or nest cell. An agent belonging to a particular colony marks each cell that it moves through with a colony-specific marker. The accumulation of such markers (with fading effects over time) facilitates the emergence of territorial movement behavior.

#### Consumer-resource ecology

When individuals move they loose biomass, but if they remain in a cell they gain biomass. Changes in biomass are computed using dynamic consumer-resource interaction processes that incorporate stress components as well [63, 64]. These dynamics account for resource growth, saturation, storage processes, and agent resource-extraction rates (which depend on the efficiency of extraction as a function of resource density, as well as the impacts of interference competition from other agents: i.e., so-called mutual interference [65, 66]).

#### Territoriality

Cells have a fading memory of the accumulated visits of all individuals from a particular colony (through the deposit of a scent-like marker each time an agent enters the cell) [67].

#### Movement

At each time-step—except when moving back to the colony at regular intervals of time—individuals move to cells, selected with given probabilities based on their relative attractiveness. This attractiveness increases with resource richness and decreases with distance, as well as with the strength of the markings of agents from foreign colonies.

#### Births and deaths

Only females reproduce with a particular probability at regular nest-cell return times (note that nothing about mating structures and male limitations are assumed), provided they are sufficiently fit (i.e., large) and not too stressed. Individuals die at a natural-mortality specified rate, plus an additional disease-induced-mortality rate during their infectious period.

#### Fission-fusion

Individuals moving into a cell that contains a colony, will join the group associated with that colony with a particular probability provided their own colony is some specified ratio larger than their current colony.

#### Force of disease transmission

Each cell presents a risk of infection to susceptible (S) agents entering that cell with a force proportional to the number of infected (I) individuals currently in the cell added to a fading value of past visits by infected individuals. This incorporation of accumulated but fading risk from past visits allows the model to deal with both direct (current I individual visits) and indirect disease transmission [68] (past visits of I individuals)

#### Epidemiology

We allow individuals to cycle through an S (susceptible), E (exposed but not yet infectious), I (infectious), V (immune), S cycle at rates respectively proportional to the risk force of disease transmission, the inverse of the latent period, the inverse of the infectious period, and the inverse of refractory immune period. Individuals in all stages are subject to a natural mortality rate, while individuals in the I stage are subject to an additional disease-induced mortality rate.

### 2.2 Technical Aspects

The mathematical equations are provided in Section 6, which are placed at the end of the manuscript because it is not necessary to go through these details before the presentation and discussion of our simulation results (those wishing to see the equations before the results should read Section 6 before starting Section 3). These equations were used to code the model using the Numerus Model Builder (NMB) platform that greatly facilitated rapid, accurate construction of the model through NMB’s hierarchical architecture and chip design. More specifically, cell equations and agent equations were first constructed as individual modules, which were then used to populate a “world” with an open-ended number of agents moving over an 18×18 hexagonal cellular array that contained 9 regularly spaced nest cells (Fig. 1). A toroidal topology was used to avoid boundary effects. The simulation period was 2000 time steps, with individuals returning every 10 time steps to their colonies to reproduce.

Some of the details on how to build such models using NMB have been provided elsewhere [26, 69], with additional novel features discussed in our Supporting Online File (SOF). Notably, the NMB platform has two independent components. The first is a set of coding tools accessed through a graphical user interface (GUI) that is used to generate a NovaScript, which is an enriched JavaScript language. This script can then be run in any suitable run-time environment, where NMB currently provides a Java runtime engine implementable in Windows, MacOS, and Linux environments. An example of a script created for our model is provided in our SOF, together with a table that facilitates moving between the code and the mathematical description in Section 6. Finally, our model is available online (see our SOF) and interested readers can run the model after downloading a free version of the NMB platform at the Numerus website (numerusinc.com).

## 3 Results

Prior experience with the consumer-resource component of the model [69, 70] and some experimentation led us to select a set of resource and consumer growth and interaction process rate parameters, as well as consumer stress and mortality process rate parameters (Tables 2 & 3) that were used to generate a baseline set of simulation results against which we could compare the agent and colony population dynamics under different movement and disease scenarios. We set up our NMB dashboard so that we could observe plots of information in real time and create a visual record of all parameter values used to produce the results of interest. An example of this output obtained during one of our simulations is provided in Fig. 2.

**Table 1.** Times to population extinct for the base line set of parameters (Table 2 except *µ* = 0.02)

| Scenario | Times to extinction for 15 runs | Average<sup>†</sup> |
| --- | --- | --- |
| No colony switching | 517, 523, 541, 818, 799, 437, 928, 437, 563, 659, 812, 475, 323, 467, 498 | 425.0 |
| Colony switching<br>($p_{\text{switch}} = 0.8$) | 310, 337, 449, 333, 454, 620, 491, 413, 701, 328, 275, 587, 364, 316, 397 | 586.5 |

<sup>†</sup>The difference is significant at the $p < 0.01$ level: two-sided Mann-Whitney U test.

**Table 2.** Model Parameter Values

| Parameter<sup>†</sup> | Symbol | Eq. | Values used |  |
| --- | --- | --- | --- | --- |
|  |  |  | Baseline | Other |
| max_rad | $K$ | 2 | 2 | fixed |
| dist_dk | $\varepsilon$ | 2 | 1/3 | 0.0, 1.0 |
| weight_res | $\phi_1$ | 5 | 1/3 | 0.0, 1.0 |
| weight_occ | $\phi_2$ | 5 | 1/3 | 0.0, 1.0 |
| max_res_ex_def | $u$ | 6 | 5.0 | fixed |
| eff_ex | $h$ | 6 | 5.0 | fixed |
| comp_ex | $q$ | 6 | 1.0 | 0.0, 10.0 |
| biomass_conv | $\kappa$ | 7 | 0.2 | fixed |
| fit_max | $a_{\alpha,4}^{\max}$ | 7 | 1.0 | fixed |
| fit_decay | $\psi$ | 7 | 0.1 | fixed |
| res_grow | $r$ | 9 | 0.2 | fixed |
| res_sat | $s$ | 9 | 20 | fixed |
| res_reservoir | $g$ | 9 | 0.1 | fixed |
| move_cost | $\nu$ | 10 | 2 | fixed |
| stress_decay | $c_z$ | 11 | 0.05 | fixed |
| res_def | $X^*$ | 12 | 0.5 | fixed |
| stress_scale | $c_m$ | 12 | 1.0 | fixed |
| mem_fade | $\eta_M$ | 14 | 0.3 | fixed |
| stress_death | $c_s$ | 15 | 1.0 | fixed |
| nat_death | $\mu_0$ | 15 | 0.01 | fixed |
| disease_death | $\mu_I$ | 16 | 0 | 0.03, 0.1, 0.2 |
| infect_fade | $\eta_I$ | 17 | 0.5 | fixed |
| disease_trans | $\beta$ | 19 | 0.0 | 0.01, 0.02, 0.03, 0.05 |
| p_switch | $p_{\text{switch}}$ | 20 | 0.0 | 0.8 |
| rho_crit | $\rho_{\text{crit}}$ | 20 | 0.3 | fixed |
| p_rep | $b_{\text{min}}$ | 21 | 0.8 | fixed |
| parent_fit | $\phi_{\text{parent}}$ | 21 | 0.65 | fixed |
| newborn_fit | $\phi_{\text{newborn}}$ | 21 | 0.35 | fixed |
| latent_inv | $\gamma_E$ | ‡ | 0.5 | fixed |
| infect_inv | $\gamma_I$ | ‡ | 0.2 | fixed |
| immunelose_inv | $\gamma_V$ | ‡ | 0.02 | fixed |

<sup>†</sup>max:=maximum, rad:=radius, dist:=distance, dk:=decay, occ:=occupy, res:=resource, ex:=extract, eff:=efficiency, comp:=competition, sat:=saturation, fit:=fitness (same as biomass in our model), def:=deficit, mem:=memory, nat:=natural, trans:=transmission, crit:=critical, rep:=reproduction, inv:=inverse
‡ See subsection on **Disease transmission**

**Table 3.** Simulation Parameter Values and Initial Conditions

| Parameter or Variable | Symbols and Values |
| --- | --- |
| Landscape dimension | $H_\gamma \times H_\delta = 18 \times 18$ |
| Simulation time | $T = 2000$ |
| colony return time | $T_f = 10$ |
| Number of colonies | $M = 9$ |
| Initial cell memory vector | $c_{\gamma,\delta,j}(0) = 0, \quad j = 1, \dots, M^\#$ |
| Initial cell resources | $c_{\gamma,\delta,M+1}(0) = U(1, 3)^{\dagger\#}$ |
| Initial cell infectiousness | $c_{\gamma,\delta,M+2}(0) = 0^\#$ |
| Initial number of agents | $N_0 = 350$ |
| Initial agent sex | $a_{\alpha,1}(0)$ is “Female” with probability $p_f = 0.6^\ddagger$ |
| Initial agent membership | $a_{\alpha,2}(0)$ randomly assigned to one of $M$ colonies$^\ddagger$ |
| Initial agent disease state | $a_{\alpha,3}(0)$ is S except for one random $E^\ddagger$ |
| Initial agent biomass/fitness | $a_{\alpha,4}(0) = U(0.15, 0.25)^{\dagger\ddagger}$ |
| Initial agent stress level | $a_{\alpha,5}(0) = U(0.15, 0.25)^{\dagger\ddagger}$ |

$^\dagger U(a,b)$ is a random drawing from the uniform distribution on $[a, b]$
$^\# \gamma = 1, \dots, 18, \delta = 1, \dots, 18$
$^\ddagger \alpha = 1, \dots, N_0$

**Fig 2.**
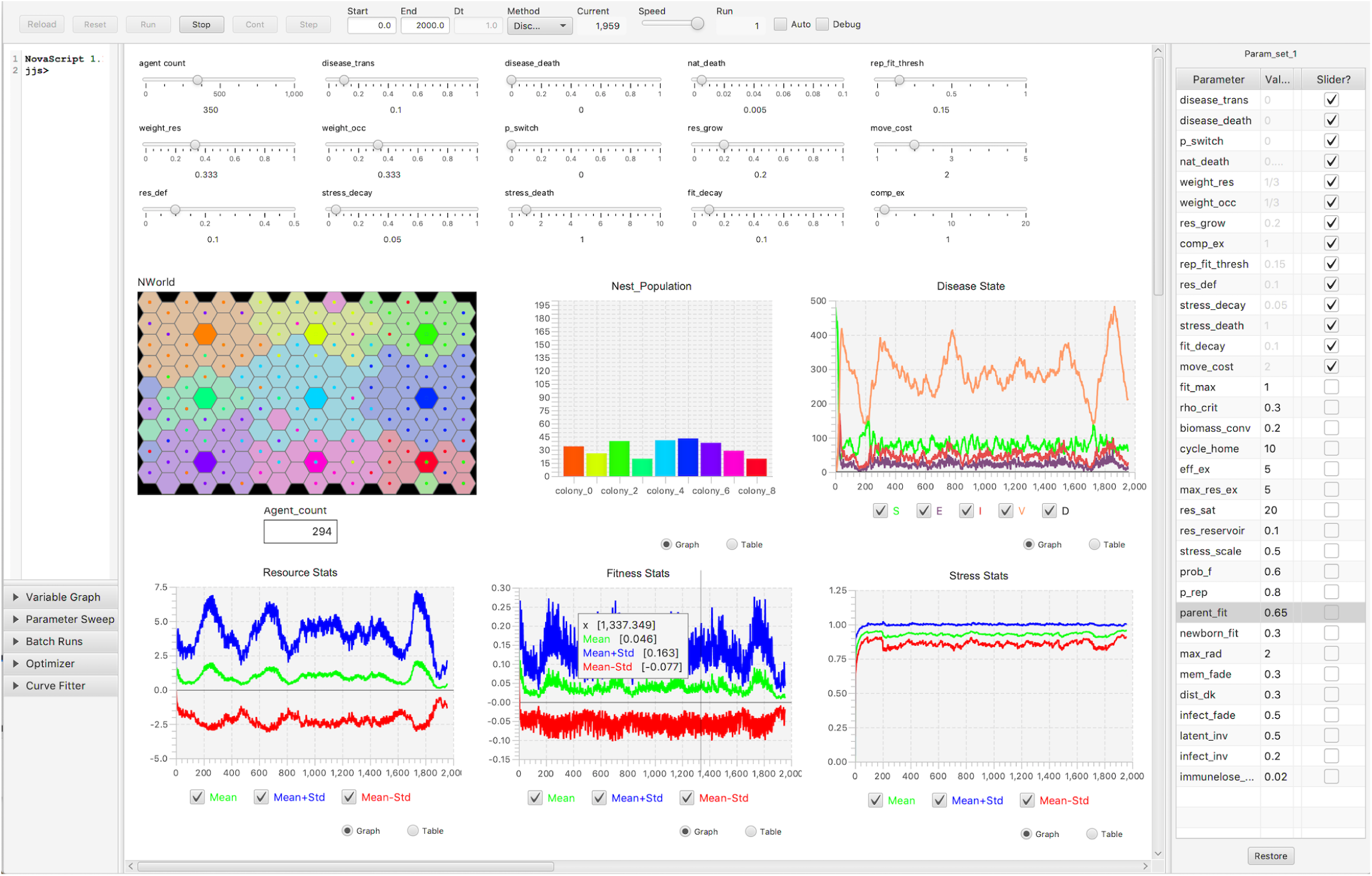
Numerus model builder (NMB) dashboard [69] with output from a simulation run are plotted up to time *t* = 1959 (see number under “current” on top strip). Parameters that can be manipulated using sliders appear above the 6 graphics panels, while a full list of parameters and values appear on the right-hand side (slider default values in light grey, other parameter values in black; also see Table 3). The top left graphic panel depicts the 18 × 18 hexagonal landscape: colony nest cells are indicated using fully saturated colors and their territory cells using corresponding partially saturated colors (note “boundary spillover” effects due to the toroidal landscape topology). The number of individuals in each colony at time *t* = 1959 are plotted in the central bar graph, using matching colony nest-cell colors. The number of agents in each disease class are plotted in the upper right-most graphics panel. Plots are also shown of the average cell resource (bottom left graphics panel), agent biomass/fitness (bottom center graphics panel) and agent stress (bottom right graphics panel) levels over time. A mouse-controlled arrow can be used to read particular values from the plotted output (bottom center graphics panel).

### 3.1 Base-line scenario: no disease, no colony switching

We ran two instances of the baseline scenario and obtained similar results with some stochastic variation across runs. A plot of the number of agents *N*_*t*_ over time (all are in the susceptible disease state) for the the two runs is provided in Fig. 3 and can are seen to be close in both period and amplitude. We note that our baseline scenario places us at a particular point in the consumer-resource growth-rate, extraction, and interaction parameter value space of our model. Different behaviors are expected at other points in this space, just as a different qualitative behaviors arise in the Lotka-Volterra prey-predator model with the type of functional response that we use in our model (i.e., Beddington-DeAngelis extended to an agent-based setting [71–73]). For example, if we decrease the mutual interference level (parameter *q* in Eq. 6) from 1 (resource extraction inefficiencies due to self and others are proportional to the total biomass of all agents in cells) to 0 (inefficiencies arise only from self) or increase it to 10 (inefficiencies increase ten fold for each competitor in the cell, presumably due to territorial conflicts) then we see a small decrease in the oscillation frequency in the former case (compare plots Fig. 3 with orange plot in Fig. 4A), but a very different type of trajectory in the strong mutual interference case (compare plots in Fig. 3 with orange plot in Fig. 4B).

**Fig 3.**
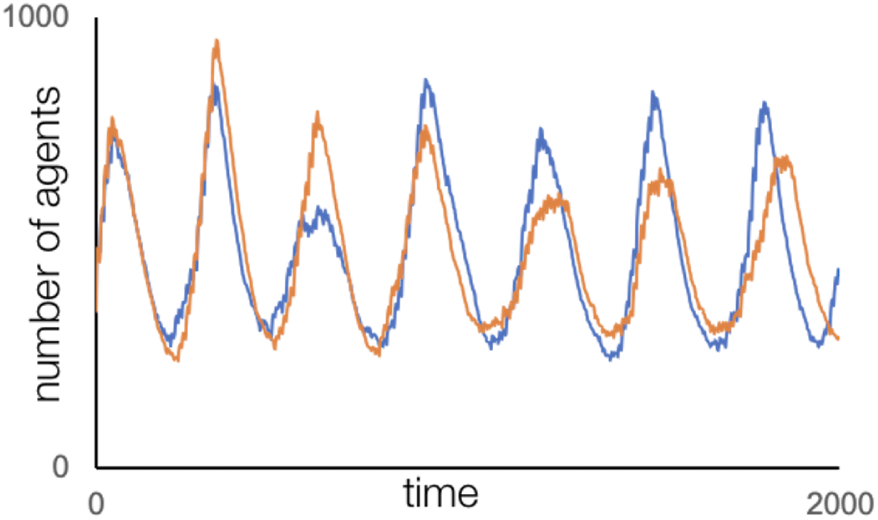
Two different instances (blue and orange trajectories) of the number of agents *N*_*t*_ obtained for the base-line simulations (no competition, no colony switching, no disease) are plotted over the 2000 step time interval.

**Fig 4.**
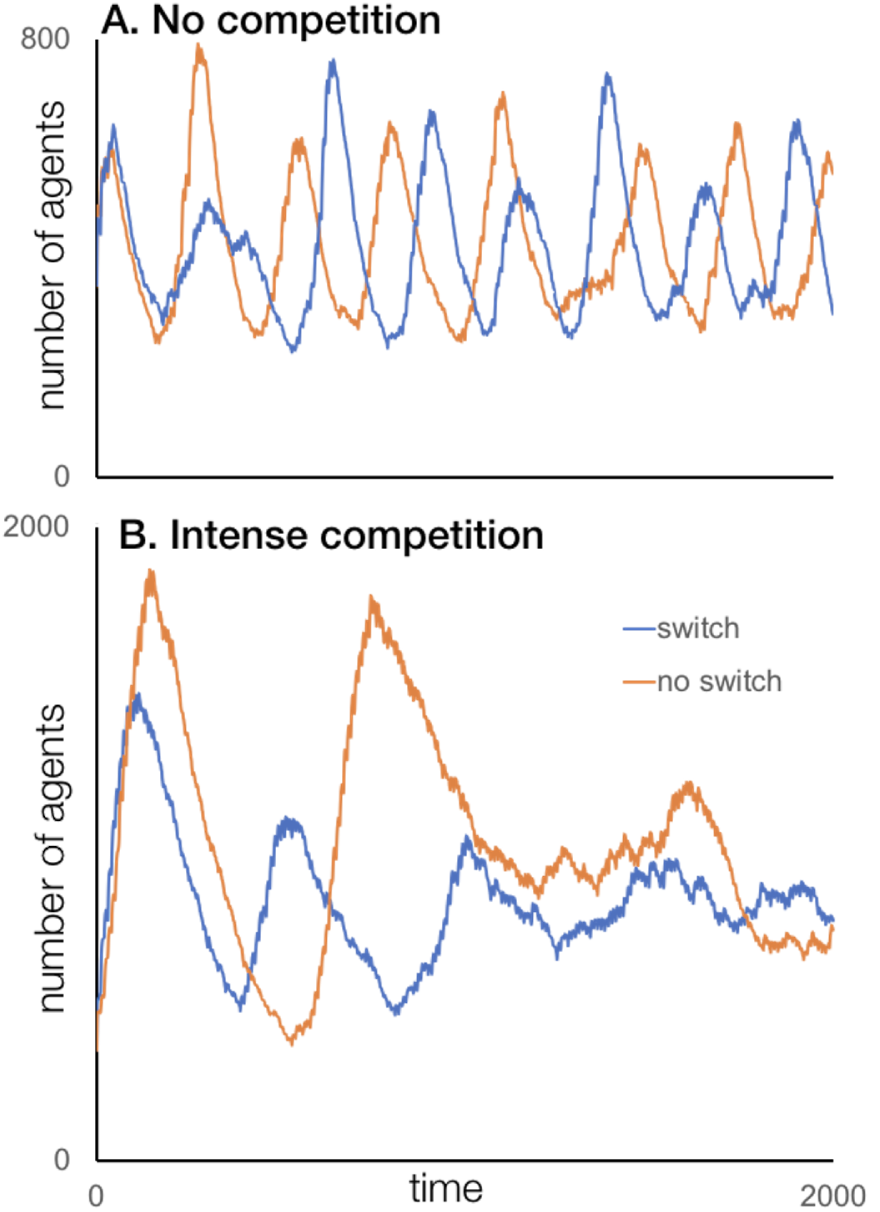
The number of agents *N*_*t*_ from the base-line simulations without (*q* = 0 implying no competition; Panel A.) and with competition (*q* = 10; Panel B.) for the no-colony-switching (*p*_switch_ = 0.0; blue plots in Panels A and B) and colony switching (*p*_switch_ = 0.8; orange plots in Panels A and B) scenarios are plotted over the 2000 step time interval. Note that the vertical scales on panels A (0 to 800) and B (0 to 2000) are different.

### 3.2 Effects of different movement strategies

We evaluated the effects of our agents moving on the total number of agents over time for the following five strategy rules (note: the rules apply to staying in the current cell or moving to one of 18 the cells within a radius of 2 hexagonal rings for all times steps except *t* mod *T*_*f*_ when all individuals must be back in their nest cells):

1. maximize available resources (*ϕ* _1_ = 1, *ϕ* _2_ = 0) (*p*_switch_ = 0.0)
2. maximize territory size (i.e., individual in colonies other than your (*ϕ* _1_ = 0, *ϕ* _2_ = 1)
3. minimize movement costs (*ϕ* _1_ = *ϕ* _2_ = 0)
4. maximize an equally weighted combination of the above three (*ϕ* _1_ = *ϕ* _2_ = 1*/*3)
5. maximize an equally weighted combination of all three, but allow individuals to switch colonies with an 80% probability when in the nest cell of a foreign colony that is no more than 30% as large as the individual’s current colony (*p*_switch_ = 0.8, *ρ*_crit_ = 0.3)

The number of agents produced by an illustrative simulation for each of movement strategies 1-4 are graphed in Fig. 5. Simulations involving movement strategy 5 were undertaken in the context of exploring the effects of mortality on population viability.

**Fig 5.**
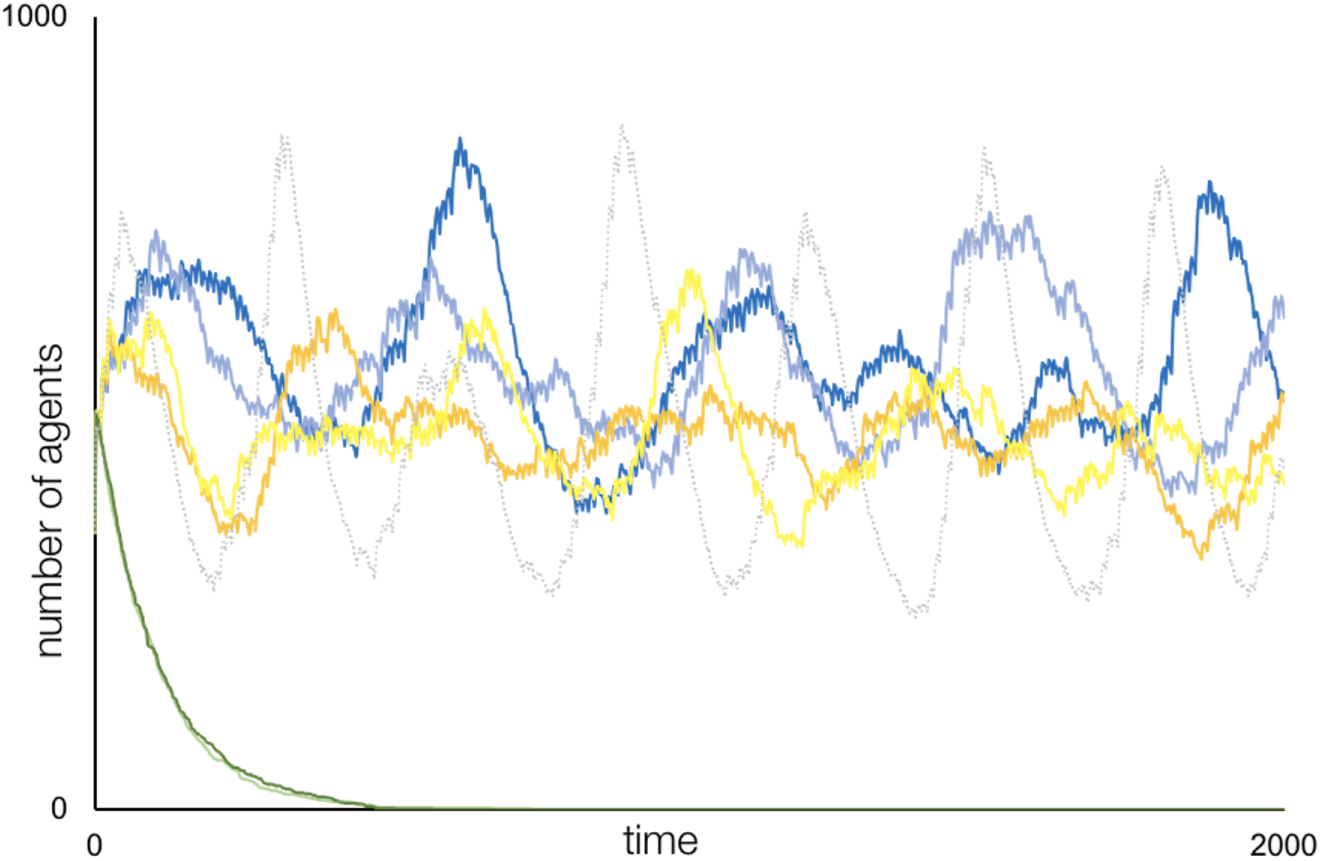
Number of agents *N*_*t*_ (i.e., the agent population trajectory) are plotted for the baseline case except for the following modifications to movement behavior. Movement is driven by: territory size maximization (blue plots, two replicates when *ϕ* _1_ = 0, *ϕ* _2_ = 1); available resource maximization (green plots, two replicates when *ϕ* _1_ = 1, *ϕ* _2_ = 0); movement cost minimization (orange and yellow plots, two replicates of *ϕ* _1_ = *ϕ* _2_ = 0); and equal weightings of above three (dotted grey plot—the same as blue plot in Fig. 3 where *ϕ* _1_ = *ϕ* _2_ = 1*/*3).

### 3.3 Mortality, colony switching and extinction

To illustrate the impact of mortality rates on population viability, we ran our baseline scenario 15 times with the natural mortality increased from *µ* = 0.005 to *µ* = 0.02. The times at which the population was extirpated in these 15 runs is given in the first row of Table 1. In the second row of Table 1, we report the result for the case where *µ* = 0.02; but now individuals are allowed to switch colonies with a probability of 0.8 (i.e. *p*_switch_ = 0.8) whenever they enter a nest cell of another colony that is no more than 30% as large as their own colony (i.e. *ρ*_crit_ = 0.3).

### 3.4 Pathogen transmission

The number of infected individuals in a population, after seeding the population with an exposed (or infectious) individual under non-zero pathogen transmission conditions, follows a U- or J-shaped bimodal distribution [74, 75] (Fig. 6): the left-hand mode represents failed or short-lived/stuttering outbreaks [76], while the right-hand mode represent successful outbreaks (i.e., the number of infected individuals rises to a peak and then falls to zero)—unless endemic conditions ensue (i.e., the presence of infected individuals persists over time) [48]. In the latter case, the bimodal distribution is essentially a long-tailed distribution, given that all stochastic population processes ultimately go to zero, if zero is an absorbing state (which it is in our model since no new infectious individuals are recruited from outside of our metapopulation) [77] (Fig. 6). Our model is initially seeded for potential disease outbreaks in that one of the agents at simulation initialization is in disease state E (exposed), while the remaining 349 are in disease state S (susceptible). To set up potential pathogen transmission outbreak scenarios requires that the transmission parameter be positive (i.e. *β* > 0). To explore the nature of these bimodal distributions for our model, we simulated 15 instances each of disease dynamics for the cases *β* = 0.01, 0.2, and 0.03. The amount of time it took for the number of exposed and infected individuals to fall to zero in each of these cases is represented in the histograms plotted in Fig. 6A. The agent dynamics in the longest-lasting of these 45 runs is illustrated in Fig. 6B. In these simulations, we note that the only effect of pathogens on host-population dynamics is through the assumption that individuals in disease states E and I do not reproduce, and not through disease-induced mortality (considered next).

**Fig 6.**
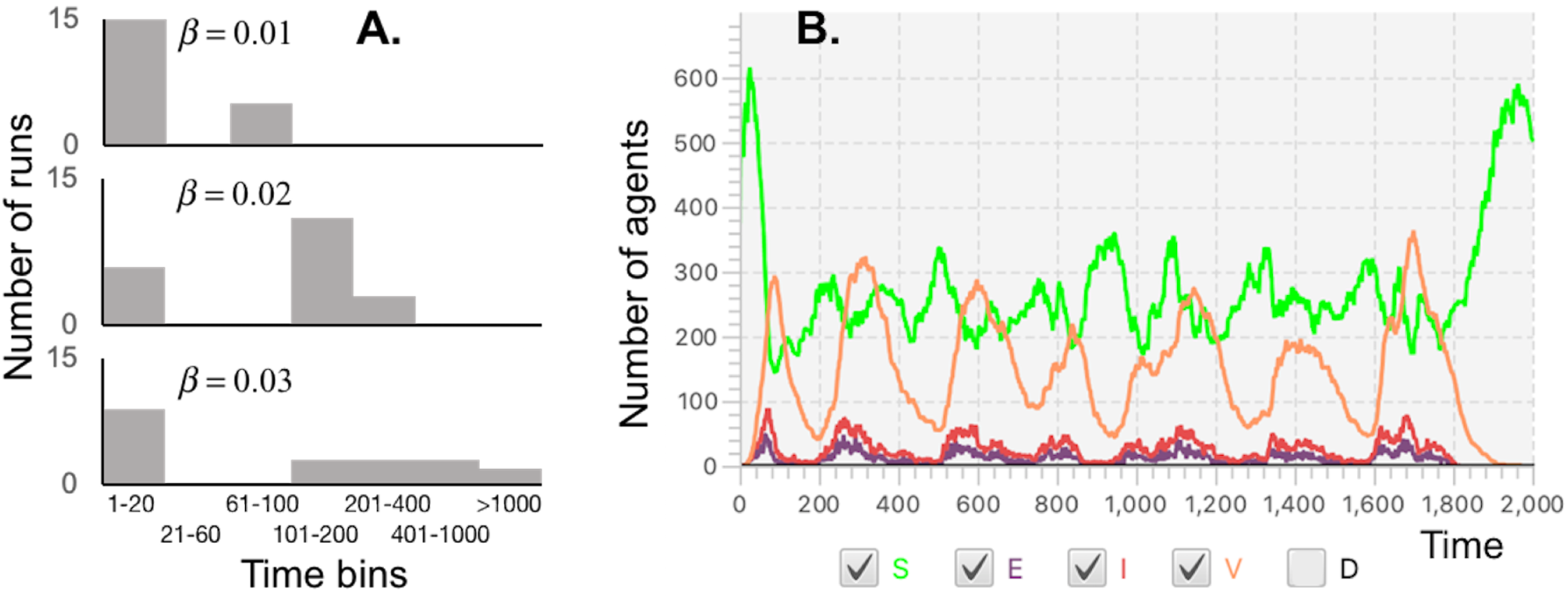
Panel A. Times taken for all infected (I) and exposed (E) individuals to be extirpated by natural mortality (disease mortality rates are zero in these simulations) are plotted as a histogram for 15 replicate simulations for each for the three transmission parameter cases *β* = 0.01, 0.2, and 0.03, as labelled (note: time scale is log-like). Panel B. Plots of the number of individuals in disease states S, E, I, and V (immune) for Run 3 of the case *β* = 0.03, where final extirpation of the epidemic occurred at time *t* = 1816 (note that number of dead individuals in state *D* was not plotted since it is an integrated—and hence steadily increasing—rather than an instantaneous value of time).

### 3.5 Disease virulence

In standard SIR epidemic models, the virulence of a pathogen is assumed to be captured by the size of the disease-induced mortality rate parameter [14]. In our model this parameter is *µ*_I_ (Table 2). We investigated the effects of increasing the value of this virulence parameter in the context of a pathogen transmission rate of *β* = 0.05 (Table 2) that leads to outbreaks that generally persist over our 2000 time step interval (cf. Fig. 6 where for *β* = 0.03 14 runs led to extirpation of disease within 1000 time steps and one at time *t* = 1816). In this case, the number of infectious individuals remained positive for the full simulation interval of 2000 time steps (Fig. 7). As virulence levels increased, the number of infectious individuals decreased and the total number of susceptible individuals increased. The highest virulence level, the number of infected individuals kept being driven down to almost zero, at which which point the number of susceptible individuals increases only to be driven down at the next outburst in the number of infectious individuals.

**Fig 7.**
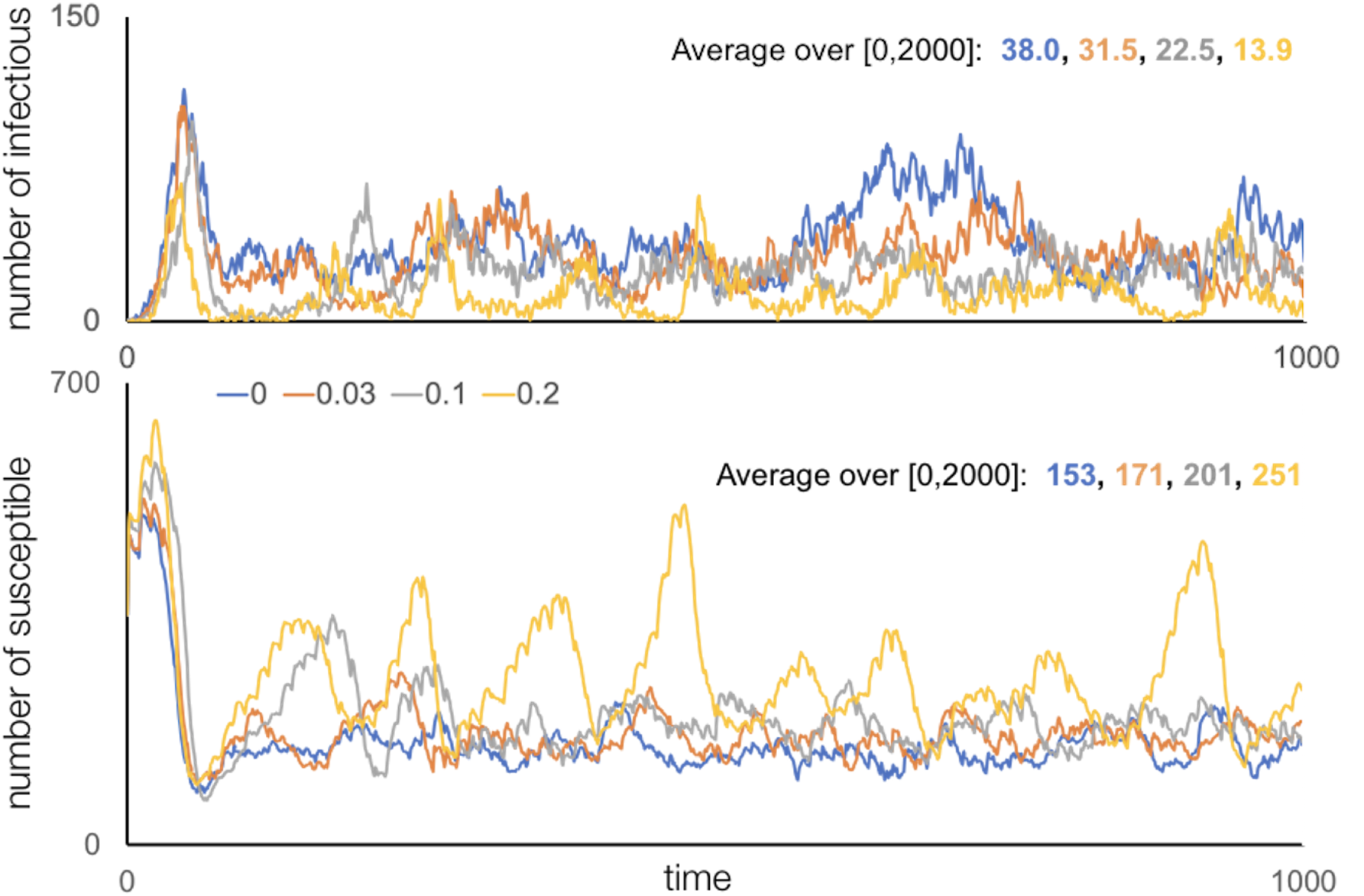
The number of infectious (I: top) and susceptible (S: bottom) agents in four simulations over the interval [0,2000] are plotted here over the first 1000 time steps only (for purposes of clarity) for the baseline set of parameter values, except *β* = 0.05 and *µ*_I_ = 0.0 (blue), 0.03 (orange), 0.1 (grey) and 0.2 (yellow). The average values were calculated over the full 2000 time steps

## 4 Discussion

The results we obtained are meant to provide exemplars of how simulations can be designed to develop quantitative narratives that have the potential to contribute to the growing body of knowledge that we call “Ecological Epistemology.” The model, as constructed, provides a tool for consumer-resource interactions to be explored in the context of individuals moving over a landscape and competing locally (i.e. in cells) to extract a renewable resource, where the extraction function has a Beddington-DeAngelis mutual interference form [65, 66, 73]. This consumer-resource interaction component has been extensively explored in the context of aggregated population level models [72, 78–80] that have been extended to include diffusion [81] and viral disease [82–84] as well.

Quantitative narratives arising from population level analyses of consumer-resource (which includes prey-predator, plant-herbivore and host-parasite/pathogen, etc.) interactions can be traced back to the work of Lotka and Volterra in the 1920’s [85] on prey-predator systems in fish populations. They articulated the principle that such interactions invariably oscillate “for some time” after being perturbed from a coexistence equilibrium, or they may support sustained oscillations “for all time.” The time concepts are in inverted commas because they relate to results obtained from deterministic models while, in effect, all population processes exhibit some level of stochasticity: as such, they will inevitably go extinct [77]. Lotka and Volterra’s work also demonstrated that density-dependent resource growth, leading to resource saturation at some carrying capacity in the absence of consumers, has the effect of dampening oscillations over time. In addition, the more recent work on systems with Bedding-DeAngelis type functional responses reveals that mutual interference in predators, as they compete to exploit resources, also has a dampening effect on population oscillations. [72].

One way to include spatial components in consumer-resource interactions is to extend their ordinary differential equation formulations to partial differential equation diffusion settings [72, 81]. Mathematical analyses in such settings are particular challenging and, even in the simplest cases, are limited to identifying whether such populations produce spatial waves or settle down to some kind of distribution, with the solution very much dependent on assumptions about the population behavior at its spatial boundary (e.g., being reflected by the boundary or migrating beyond the boundary). As soon as population behavior takes on complexities that lead to various kinds of nonlinearities in equations expressing those complexities, either numerical solutions are needed or population behavior can simulated using agent-based models [30]. Agent-based models are particularly useful when considering stochastic aspects of consumer-resource interaction processes.

Agent-based versions of prey-predator processes have been built using the Netlogo [86] and Starlogo [87] modeling platforms. The first of these modeled predation at a “fast” minute-by-minute time scale. Predation events were calculated based on correlated random walks and accounting for predator perceptual ranges and prey refugia. Predation events were then accumulated along with births and deaths into a Lotka-Volterra-type population process iterated at a “slow” daily time scale. This study found *inter-alia* that increasing predator efficiency by either increasing their perceptual range or decreasing the number of prey refugia resulted in a coexistence region—an equilibrium around which the prey and predatory populations oscillated—decreased for both prey and predator, thereby increasing the risk of extinction. As in our model, the second study included the interaction of both competition and predation. The quantitative narrative the emerged for this study [87], quoting from its abstract, was:

> *In a low-production environment, the smaller and faster consumer out-competes the larger and slower one, but in a high production environment the larger and slower consumer survives. Predation hastens the extinction of one of the consumers, but niche partitioning of the consumers increases both the coexistence of consumers and the number of predators. Predators with medium efficiency are able to coexist in the system longer and in larger numbers*.

One thread of the quantitative narrative that emerges from simulations of our model is that competition causes the period of prey-predator oscillations to increase and be dampened more rapidly, with this increase and dampening accentuated as colony mixing increases. A second thread is that a “grass is greener syndrome” (i.e., movement decisions are based purely on resource considerations) leads to increased stress in individuals and, as a possible result, to population collapse because individuals are not taking into account the cost of moving (green plots in Fig. 5). This collapse (green plots Fig. 5) is mitigated by territoriality considerations, since these curtail movement and allow more time for resource extraction (blue plots in Fig. 5); but movement based on territoriality-only considerations leads to dampening, albeit less so than movement-cost-only considerations (compare yellow with blue plots in Fig. 5). Notably, a movement strategy that accounts for all three factors provides the most regular, albeit oscillatory, population trajectories (dotted grey plot in Fig. 5).

A second thread of the quantitative narrative that emerges from our simulations relates to the effects of disease on population dynamics, for which our model provides an appropriate formulation to address the questions of interest. One of these effects is the impact of increasing pathogen transmission rates on the maintenance of an endemic infection. No subclass of a population (e.g., infectious individuals or even the population as a whole) can persist forever—both are bounded stochastic population processes that are known to go extinct in a finite amount of time [74, 75]. This is amply demonstrated in our simulations that consider the impacts of increasing pathogen transmission rates. In Fig. 6A, we verify that our infection process produces the expected U-shaped bimodal distributions of infectious individuals, though the breadth of the distribution associated with the outbreak mode (as opposed to an early-fade-out mode) increases with *β*. Indeed, for *β* = 0.05, once an outbreak occurs, it generally lasts longer then 1000 time steps, as illustrated in Fig. 7. This remains true as the virulence of the pathogen increases from *µ*_I_ = 0 (blue plots in Fig. 7) through *µ*_I_ = 0.03 (orange plots in Fig. 7) and *µ*_ID_ = 0.1 (grey plots in Fig. 7) to *µ*_I_ = 0.2 (yellow plots in Fig. 7). What is clear from these simulations is that as virulence increases, the number of infectious individuals decreases, though in our highest virulence case we obtain a sequence of fade outs to near extirpation, followed by outbreaks that are then contained by the high mortality rates of infectious individuals and the switching of those individuals surviving the infection to a state of temporary immunity. This immunity once lost creates, along with births, a rising group of susceptible individuals. Such outbreaks are similar to what we see in seasonal influenza—although, in this case, loss of immunity is hastened by the existence of different influenza strains and mutations in the influenza virus antigens [88]).

## 5 Conclusion

A deeper exploration of the various issues examined in this paper, as well as comprehensive investigations of the many other questions that could be asked along the lines of those we posed, would require much more extensive simulations than has been reported here. This is particularly true of simulations involving evaluations of times-to-extinction of populations [89], consequences of different movement strategies on the formation and stability of territories [67], extirpation of disease outbreaks [76] because of the stochastic nature of the processes involved. In the context of developing quantitative narratives, a point is reached where additional simulations will not add to the qualitative character of the information obtained and the increased precision that comes from additional runs is not useful. In other words, there is little value in being more precise than the level of the errors associated with the fit of models to data.

The model we present here has the potential to address a range of questions relating to the ecology of consumer-resource interactions played out over spatially-structured landscapes and impacted by movement and disease processes. Our model can also be elaborated to address questions that include, for example, mating and genetic structures, multispecies situations, environmental factors that exhibit seasonal variation, and landscape structures, including individuals moving over real landscapes. Numerus Model Builder has the facility to import landscape information from GIS databases that can be easily extended to metapopulation settings [90]: thus our model can be readily extended to ask questions about spatially-structured populations on real landscapes.

Of course, considerable interest exists in fitting models to data, especially in the context of wildlife management [91] and predicting the behavior of ecological systems that, for example, include a sociological component [92]. Agent-based models provide the best, if not the only, approach to addressing many of the more complex questions relating to the management of real socio-ecological systems and to the advancement of ecological understanding through its application to developing quantitative narratives in ecology (Fig. 1). Thus, in the future, we can expect agent-based models to increasingly be used to develop quantitative narratives as a way of advancing ecological epistemology.

## 6 Technical Methods

### 6.1 Technical Framework

In this technical presentation all `TEXT` in this font refers to an actual Numerus Model Builder built-in function, but its designation is sufficiently clear so that the algorithm can be implemented in other programming languages.

1. The population initially consists of *N*_0_ agents; but over time, due to births and deaths this number will change and be designated *N*_*t*_.
2. The landscape is a pixelated *H*_*γ*_ × *H*_*δ*_ hexagonal cellular array with a doughnut topology. The index pair *γ, δ* refers to a particular cell in this array. To emphasize that these indices refer to one entity (a particular cell), we tie them together using a hat notation: viz., 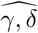. This is particularly useful when we use multiple subscripts to refer to cells in the neighborhood of the cell 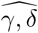.
3. *M << N*_0_ of the *H*_*γ*_ × *H*_*δ*_ cells are identified as nest cells because they are the home cell of, at most, one uniquely identified colony.
4. Initially, every one of the *N*_0_ agent is assigned to a nest cell at random. Thus, initially, there are an average of *M/N*_0_ individuals per colony. The *m*^*th*^ of these colonies is referred to as *A*_*m*_, *i* = 1, …, *M*.
5. The model has a *T*_*f*_ -unit fast-time (within) cycle that begins with all individuals at home (in their nest cells) at REMAINDER(TIME()/*T*_*f*_)=0, while INTEGER(TIME()/*T*_*f*_) is the number of cycles that have been completed.
6. At each tick of the clock except the last—i.e, for time progressing from REMAINDER(TIME()/*T*_*f*_) =*τ* to *τ* + 1 for *τ* = 0, …, *T*_*f*_ − 1, an individual can stay in its current cell or move to any of a set of cells in set of cells called BLOCK (see Eq.1 below). However, when REMAINDER(TIME()/*T*_*f*_) =0 the individual moves (jumping, if necessary) back in its home cell.
7. At each tick of the clock, except for the last within-cycle tick, when individuals return home, individuals move to increase their potential to extract resources from cells or claim new cells for their colony according to the priorities laid out in the movement rules section below.
8. Cells have a state vector of dimension *M* + 2, indexed according to the position of the cell in the array using the row index *γ* = 1, …, *H*_*γ*_ and column index *δ* = 1, …, *H*_*δ*_. The first *M* elements of each cell vector 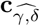 contain “fading memories” of the “occupation history” of individuals from each of the *M* colonies that have occupied this cell over time. In particular, if an individual from colony *j* occupies the *γδ* cell at time *t*, then the value of the *j*^*th*^ element 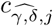 is increased by 1, but the value itself declines geometrically over time at a rate *η*_M_. The *M* + 1^*th*^ element of 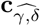, denoted by 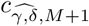, is a real non-negative number that reflects this cells current resource value. Finally, the element 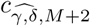 contains the infectiousness value of the cell.
9. Individual agents have a state vector **a**_*α*_, *α* = 1, …, *N*_*t*_. The elements of **a**_*α*_ are as follows:
  a. *a*_*α*,1_ is a positive integer value that designates the type of individual with the number of possible types reflecting the number of species of interest, their sex, and any other category designator (e.g., if there are two sexes for each of two species, this number may be 1, 2, 3 or 4).
  b. *a*_*α*,2_ is an positive integer value *m* = 1, …, *M* that identifies the colony membership of individual *α*.
  c. *a*_*α*,3_: this is a positive integer value (or named state) that identifies the disease state of individual *α*. This can be relatively simple—such as disease states (with a numeric equivalents) S (0), E (1), I (2), V (3). In more detailed cases, we might want to increase the number of categories so the value *a*_*α*,3_ might reflect the number of time periods since the individual was infected. The probability of being in state E, I, or V (and even back to S) could then depend on the value of *a*_*α*,3_, conditioned on the value of other elements of **a**_*α*_ if needs be.
  d. *a*_*α,ℓ*_, *ℓ* = 4, 5, … may be continuous state variables that measure the relative size of individuals, their state of hunger, their state of emaciation when food is in short supply, and so on. Here we only use two such variables and refer to *a*_*α*,4_ interchangeably as the fitness or biomass variable and *a*_*α*,5_ as the stress variable.

### 6.2 Movement rules

In this section, and in NovaScript, an extension of the JavaScript programming language used to construct Numerus Model Builder, the local cell occupied by an individual is referred to as its `RING(0)` cell. Its six immediate neighbors are its `RING(1)` cells, while all cells that it can reach in a minimum of *k* moves without jumping over any cells is its `RING` (*k*).

Recall that 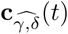 represents the state of the cell occupied by individual *α* at time *t*, where 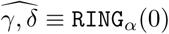 is one way of referring to the cell currently occupied by individual *α*. With this in mind, we use 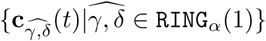 to refer to the set of state vectors of the 6 cells surrounding the current location of individual *α*. Before moving on, we stress that the designators 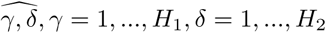 are global locators of cells, while `RING`_*α*_(*r*) is a local locator of all cells a distance *r* from the current cell occupied by individual *α*.

At the start of each time interval [*t, t* + 1], we first determine whether or not individual *α* = 1, …, *N* remains in its current cell or moves to some other cell in the set `BLOCK`_*α*_(*r*), where *r* is some pre-selected positive integer that determines the furthest distance any single move can be:

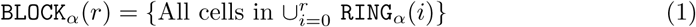

The total number of cells in this set is *K* = 1 + 6(1 + 2 + … + *r*) = 1 + 3*r* + 3*r*^2^ (one center cell plus each hexagonal ring centered on this center cell ring has 6 cells more then the its immediate inscribed ring: i.e., 1+6+12+…+6(r-1)+6r cells). We use the notation 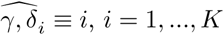 to index a renumbering of cells in `BLOCK`_*α*_(*r*). One renumbering scheme, for example, is to label the center cell 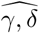, occupied by individual *α*, as 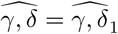, the cell to its immediate west as 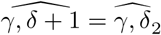, then going counter-clockwise until the last of the cells in `RING`_*α*_(1) has been labeled 7 (i.e., 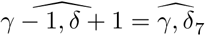 and moving on to the west most cell in RING_*α*_(2) as 8 and so on until all cells in `RING`_*α*_(*r*) have been labeled.

The decision for agent *α* belonging colony *ℓ* to stay in its current cell or move to some other cell in `BLOCK`_*α*_(*r*) is determined using the agent movement rule below that relies on the following three measures:

#### Distance measure

*Closer is better*

For some decay parameter *ε* > 0, distance radius *v*_*i*_, and maximum distance radius *K*

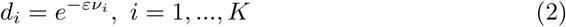

represent the distance between the center cell and the *i*^*th*^ cell in `BLOCK`_*α*_ when this cell is in `RING`_*α*_(*v*_*i*_).

#### Comparative resource measure

*Prefer cells with more resources*

Under our renumbering 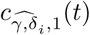 is the resource value of the *i*^*th*^ cell in `BLOCK`_*α*_ at time. Use this to calculate for *i* = 1, *K*,

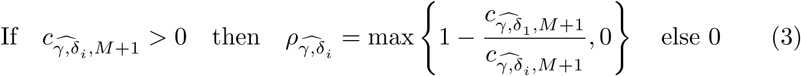

#### Comparative occupation-history measure

*Prefer cells less used by other colonies*

Under our renumbering 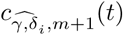 is cell *i*’s 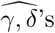 memory state with regard to the occupation history of agents from colonies *m* = 1, …, *M* in this cell. For cells *i* = 2, …, *K*, this quantity is based on a comparison of the largest occupation history value (in cell *i*) of 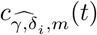 for all *m* = 1, …, *M* in comparison with the occupation history value of agent *α* in cell *i*. Recalling that agent *α* is from colony *ℓ*) then, for *i* = 1, *K*,

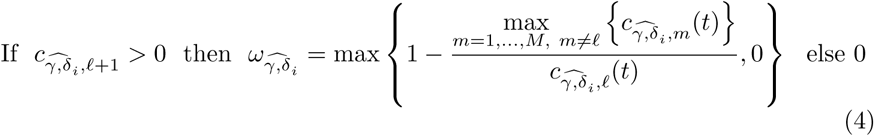

#### Agent movement rule

*Stochastic trade-off*

For weighting parameters 0 ≤ *ϕ* _1_ ≤ 0 and 0 ≤ *ϕ* _2_ ≤ 1 − *ϕ* _1_—which represent trade-offs among the relative values of movement distance, resources of cells, and past use of cells—define the cell attractiveness values *p*_i_, *i* = 1, …, *K* as

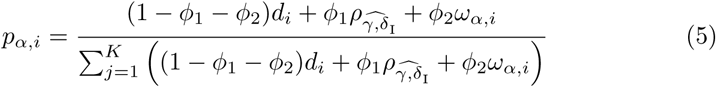

Now choose among the cells by applying the single drawing multinomial distribution: MULTINOMIAL[1;*p*_*α*,1_, …, *p*_*α,K*_ ]

### 6.3 Ecological Dynamics

First, we determine for all cells which individuals occupying each cell stay or move, and then, we compute changes in the resource states of the cells, as well as the fitness that individuals gain or lose from consuming resources due to movement activity.

#### Movement *Who moves?*

Assume that there are 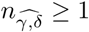 in cell 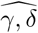 individuals at times *t* that are not divisible by *T*_*f*_. Then for each of these individuals, a multinomial drawing based on Eq. 5 is implemented to see if an individual stays or moves; and, if it moves, to which cell it moves.

#### STAY dynamics *Cell resources and individual fitness updating*

If 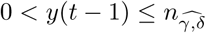 individuals stay in cell 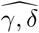 over the time interval[*t* − 1, *t*] (i.e., these individuals were in cell 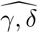 at time *t* − 1 and did not move out at time *t*), then the individual and cell states are updated using the following quantity defined in terms of the maximum resource extraction rate parameter *u* > 0, extraction-efficiency parameter *h* ≥ 0, and competition parameter *q* > 0 (see [69, 70], taking into account that *y*(*t* − 1) − 1 is the number of individuals in each cell beyond a focal individual for which the extraction expression is being formulated):

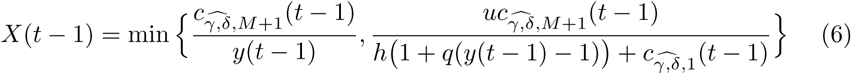

If *y* = 0 then *X*(*t*) = 0. Specifically, the individuals’ fitness states *a*_*α*,4_(*t* − 1), *α* = 1, *N*, are updated for a maximum fitness value 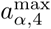, a fitness decay factor *ψ*, and a biomass conversion parameter 0 *< κ <* 1 using

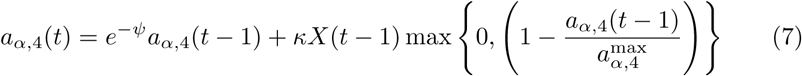

and the cell 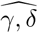 resource state for resource-intrinsic-growth-rate parameter *r* > 0, reservoir parameter *g* ≥ 0 and saturation parameter *s* > 0 is updated, by first defining

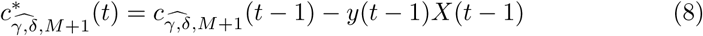

and then using the equation

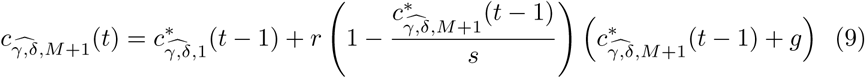

#### MOVE dynamics *Individual fitness updating*

If an individual *α* moves out of a cell at time *t* − 1, then the individual’s fitness (or biomass) decays at a rate *vψ* > 0, where *ψ* is the decay rate factor introduced in the STAY dynamics section and *v* > 0 is the fitness cost of moving relative to not moving; i.e.,

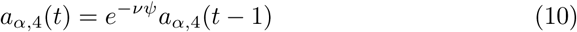

#### Stress dynamics *Individual not meeting daily requirements*

We model stress dynamics using a variables *a*_*α*,5_(*t*) for the *a* = 1, …, *N* agents. We note that *a*_*α*,5_(*t*) ≥ 0, such that *a*_*α*,5_(*t*) = 0 implies the individual is unstressed with stress increasing as *a*_*α*,5_(*t*) increases. We assume that without additional stress input, the *a*_*α*,5_(*t*) relaxes back to zero at a rate *c*_*z*_. However, if an individual is subject to an additional stress *z*_*α*_(*t*) over the interval [*t, t* + 1] then its new stress level at time *t* will be

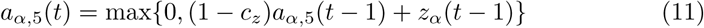

To calculate the amount of stress an individual is subject to in each time interval, we assume that when it moves (i.e. does not take in any resources) then it has a resource deficit of 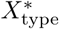 (i.e., a set of input parameter values subscripted by type, where “type” indicates that these deficits may vary by species, sex, or age, etc.). In this case we assume, dropping the type designation (since we make this parameter the same for both sexes, all disease states, and we have no incorporated age structure), that

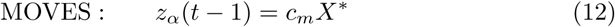

where *c*_*m*_ > 0 is a conversion constant that depends on the units used to measure resources. On the other hand, if an individual gets to stay in the cell over time [*t, t* + 1], and there are *y* individuals (including itself) that stay to exploit resources in the cell during this time interval, the amount of resource that the individual extracts, from Eq. 6, is *X*(*t* − 1)*/y*. We now assume that the extent to which this value falls short of the resource deficit value *X*\* is the amount of stress incurred over [*t, t* + 1] multiplied by the conversion constant: i.e.,

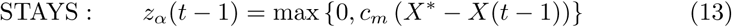

Note the the above covers all individuals that stay whether or not they reproduce.

#### Cell memory dynamics *Which colony dominates particular cells*

The quantities 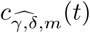, *m* = 1, …, *M* which contain a fading memory of how many individuals from colony *m* have visited the cell over time, are updated using *y*_*m*_(*t*) to represent the number of individuals from colony *m* that are in cell 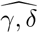 at time *t* and a memory fade rate constant *η*_M_ > 0. The updating equation for *m* = 1, …, *M* is:

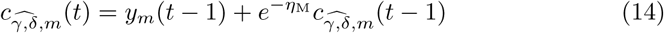

### 6.4 Epidemiology and Deaths

The epidemiological component is essentially an S (susceptible), E (exposed but not yet infectious), I (infectious), V (recovered with immunity) and D (dead) model [26]. We use the value of *a*_*α*,2_ to denote this state and use the following rules to compute whether or not individuals in states S, E, I, and V respectively transition to states E, I, V, S and D in terms of “transition rate parameters” *γ*_E_ > 0, *γ*_I_ > 0, *γ*_V_ > 0, *γ*_S_ > 0 and *γ*_D_ ≥ 0. We assume a basic death rate over the interval [*t, t* + 1) *µ*_0_ > 0 that increases with stress at rate that is proportional to the stress level *a*_*α*,5_(*t*), with a constant of proportionality given by *c*_*s*_ > 0. In this case:

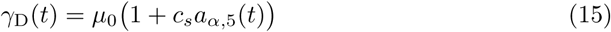

for all individuals in diseases states S, E, I and V. For individuals in disease state I, we assume an additional disease induce mortality rate 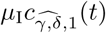 (where for relatively high mortality diseases *µ*_I_ >> *µ*_0_) so that for individuals in I, where we add the index I to distinguish it from the rate we have in Eq. 15

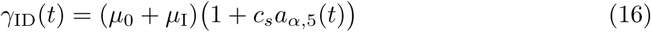

#### Cell Infectiousness *Pathogen density in each cell*

If *y*_I_(*t*) is the number of infected individuals in cell 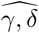 at time *t* and *η*_I_ ≥ 0 is an infectiousness fade rate (*η*_I_ close to 0 implies that indirect transmission of pathogens is of considerable importance while *η*_I_ → ∞ implies that direct contact becomes increasingly important for transmission (for a discussion on direct versus indirect transmission, see [34]), then:

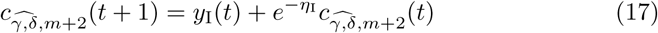

#### Transmission or death *Who gets infected?*

A susceptible individual that spends the period [*t, t* + 1) in cell 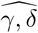 will have probability of getting infected at a rate proportional 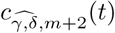, where the constant of proportionality is *β* > 0. Thus the per-capita rate or force of infection is:

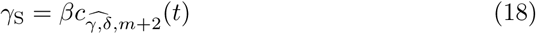

If susceptible, individuals also have a probability of dying over the same time interval at a rate (or force of risk) that is proportional to their stress level *a*_*α*,5_(*t*) then, in essences, we are assuming that the transition rate out of then using a competing rates approach [26] we obtain for *γ*_S_ and *γ*_D_ given by Eqs. 18 and 15

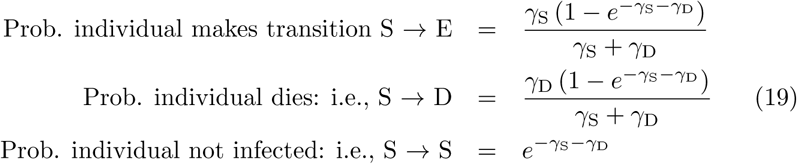

#### Disease transitions *what is an individuals next disease state?*

The transitions from states E to I or D and V back to S or D follows the same formulations of Eqs. 19, except we replace *γ*_S_ with *γ*_E_ and *γ*_V_ in each of the two cases respectively. The transition from state I to V or D, also follows the same formulation of Eqs. 19 except now we replace *γ*_S_ with some *γ*_E_ > 0, and we replace *γ*_D_ with *γ*_ID_ = defined by Eq. 16

### 6.5 Colony Switching

The number of individuals in each colony essentially constitutes a bounded stochastic walk on the non-negative integers. The upper bound is imposed by feedbacks that relate to the effects of increasing death rates and decreasing birth rates through reductions in fitness and increases in stress levels due to competition for limited resources. Such walks are known to ultimately hit zero [74, 75], which is an absorbing state for the number of individuals in each colony, unless colonies themselves can be fueled through emigration.

There are several ways to deal with emigration. A relatively simple approach is for an individual to make a decision to join a colony, when it enters a colony cell and conditions are favorable compared to the individual’s current state. For example, the individual on entering a cell can compare its own current fitness and stress levels with those of the individuals residing in the cell being entered.

Here is a very simple approach that we employ in our simulation, based on the idea that if a colony consists of a very few individuals, then competition for resources is less intense in the neighborhood of this colony. In rule articulated below, #*m* refers to the number of individuals in colony *m*:

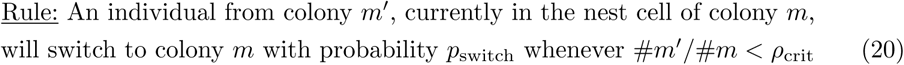

### 6.6 Reproduction

All individuals in colonies that do not move have an opportunity to reproduce at that start of each new fast cycle (i.e., when *t* mod *T*_*f*_ = 0). In particular, any individual who is in disease state S or V, and whose fitness value exceeds *b*_min_ will not move in the next time step, but will instead give rise to a new individual. The fitness/biomass value of this reproducing individual will drop (parent) and a newborn will be created with updated fitnesses given by

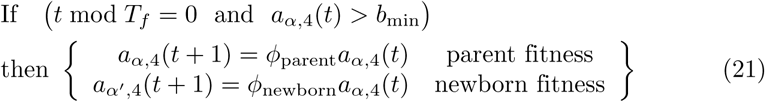

Note that the newly reproduced individual, *α*′, will belong to the same colony. In addition, we must have *ϕ* _parent_ + *ϕ* _newborn_ *<* 1, where the shortfall of this sum from 1 is a cost of reproduction.

### 6.7 Parameters and Initial Conditions

A list of all the parameters appearing in the model, the names of these parameters in our NMB code, their mathematical symbols and the equations in which they first appear, their baseline values, and other values used in non-baseline runs are given in Table 2. Values used to define the size of the simulations (size of landscape, length of runs) and the initial values used in our various simulations are given in Table 3.

## Acknowledgments

The development of Nova, a precursor to Numerus Model Builder, was supported by NSF grant CNS-0939153 to Oberlin College (PI: RS) and NSF-EEID grant 1617982 (PI: WMG). We thank John Pataki of Logical Laboratories for his considerable help and input into creating and supporting the Numerus Model Builder website http://www.numerusinc.com/.

## Authors’ contributions

WMG developed the mathematical model and wrote the first draft of the manuscript. RS coded the model and ensured its smooth running on the Numerus Model Builder platform. WMG carried out the simulations and generated the figures. KT developed the Supporting Online File, and contributed to the verification of coding and mathematical equation accuracy. All authors helped edit the the manuscript and approved the content of the submitted version.

## Supporting Online File

This file, containing details of the Numerus Model Builder construction of our model and a copy of the NovaScript file (superset of JavaScipt and implemented in a Java Run Time environment), is either attached herewith or available at a site to which this file is posted.

## Supporting Online File

### 1 Model Simulation Framework

Here we provide some of the key details not included in previous publications [1,2] regarding the construction, using the Numerus Model Builder platform, of the agent-based model presented in the main text.

Numerus Model Builder (NMB) employs *NovaScript*, which is embedded in, and extends, Javascript. The NovaScript interpreter is an extension of the ECMA 1.7 Javascript standard. The full range of the Javascript language is available to the NovaScript programmer for specifying the computations required by his or her simulation. It includes facilities for specifying simulations visually, using stock and flow diagrams (familiar to programmers of Stella and Vensim) and other visual tools unique to NMB. Simulations are scoped out using such visual schematic diagrams with equation and other procedural details entered into appropriate windows [1], or textually, via NovaScript. We will refer to the former as a diagram specification; the latter will be referred to as a code specification.

NMB is highly modularized due to: 1) the creation of dynamic agent “Capsules” that interact with their environments through input/output interfaces (more below) and 2) allowing these agents to move over a heterogeneous resource landscape. Movement can either be on a Cartesian plane or from one cell to another on a rectangular or hexagonal array of specifiable dimension [1]. Our model moves agents over hexagonal arrays of variable and dynamic resource cells and was designed to address questions regarding agent and colony populations dynamics under different movement and disease scenarios.

### 2 Model Components

Simulations are built out of objects, called Components. Among the component types are **Primitive Components, Simulators, Displays**, and **Controls**. The primitive components consist of Stocks, Variables, Sequences, Flows, and Terms. Stocks, Variables and Sequences are used to represent the state of a running simulation. The Stock component may be attached to one or more Flows. Each Flow specifies some change in the current value of the Stock. A Variable is similar, only it is not attached to any Flow; rather its change is determined by a derivative contained in the *Prime* field. Sequences are similar to variables, except that they are discrete, and change is indicated by a *Next* field, directly determining the value of the sequence in the next time interval. Flows and Terms both have *Exp* fields containing their expressions. As mentioned above, Flows affect the value on the Stocks to which they are attached. A Term is used to compute a value using current State values (in Stella, what Numerus Model Builder calls a Term is referred to as a Converter.)

This model is formulated as a set of agents moving over a cellular array. Therefore, as described in our presentation, we require 3 Capsule types: the Cell, Agent, and sirTerrM. (Refer to Sections 3.1, 3.2, and 4 for further detail on each Capsule type.)

#### 2.1 Architecture

A Numerus model is structured using elements called Simulators. Here is a description of the Simulators used in our model.

##### Simulators

The term-of-art *Simulator* is used by Numerus Model Builder to describe a container of interacting components. In a running simulation, a Simulator is set to an initial state. As the simulation progresses its next state is determined by its current state and by other simulators with which it may be communicating. The Capsule is the simplest Simulator, and may contain any of the components described in the previous section. The five other Simulator types are array-like *aggregators* comprised of Capsule elements. Each aggregator introduces new functionalities to their constituents, as described below.

##### Capsule

The role of the Capsule is to contain the components of a simulation and synchronize the progress of the computation. A Capsule holds instances of the primitive components, but may also contain other Capsules and other aggregator Simulators. The top-level construct in any model is a Capsule.

##### CellMatrix

A *CellMatrix* is a spatial array of individual Capsules, or *Cells* laid out in either a Cartesian lattice or hexagonal grid. Each Cell has a pair of coordinates and can query the CellMatrix to determine the other Cells in its neighborhood, with which it can exchange information. A CellMatrix can also provide an environment for agents to exist when combined with an AgentVector (see below).

In our model each cell contains resources (i.e., plants, fruits, seeds), can act as a nest site associated with a specified colony of agents, and can be occupied by one or more agents who either pass through in a single time period or remain for some time during which they exploit the dynamically changing (through both growth and extraction) resources.

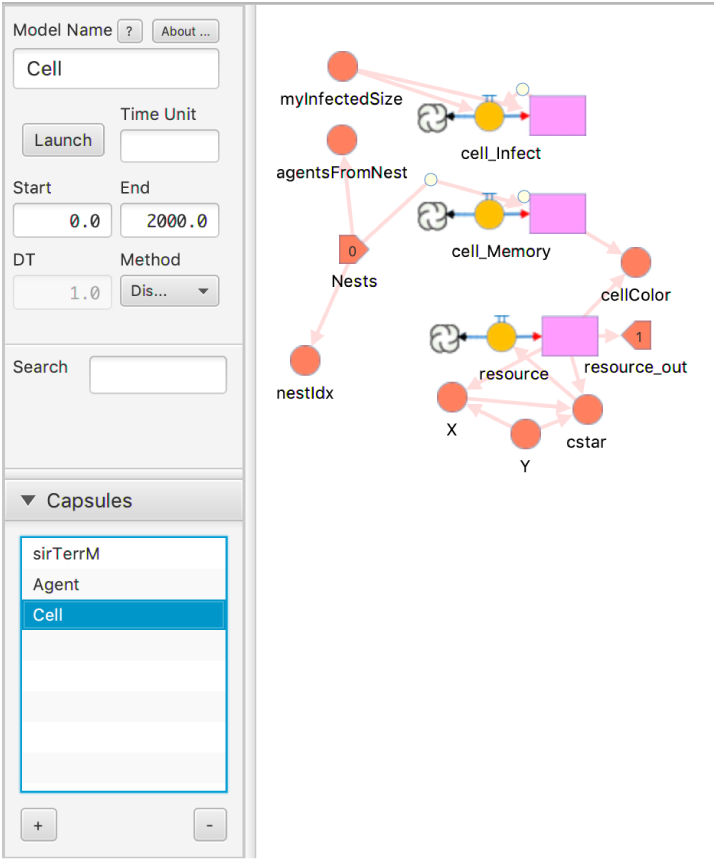

##### AgentVector

An *AgentVector* contains a dynamic set of Capsules called *Agents* that can exist on some landscape. At any point in time an Agent has a location on that landscape, which will change if the Agent decides to move. Agent motion will generally either use a toroidal topology or reflect off the boundary like a billiard ball (other boundary effects can be implemented). The AgentVector helps Agents find nearby Agents with which they may interact. New Agents may be created and existing Agents may be removed during the lifetime of the simulation.

In our model each Agent has a biomass (a surrogate measure of the agent’s individual fitness), a stress level, and a sex (determined at birth). Females may reproduce on regular returns back to their colonies. This works via the parent retaining the majority of the biomass, the offspring acquiring a minority of biomass, and some loss from the cost of the reproduction event itself. Our movement algorithm uses a toroidal topology in order to eliminate boundary effects.

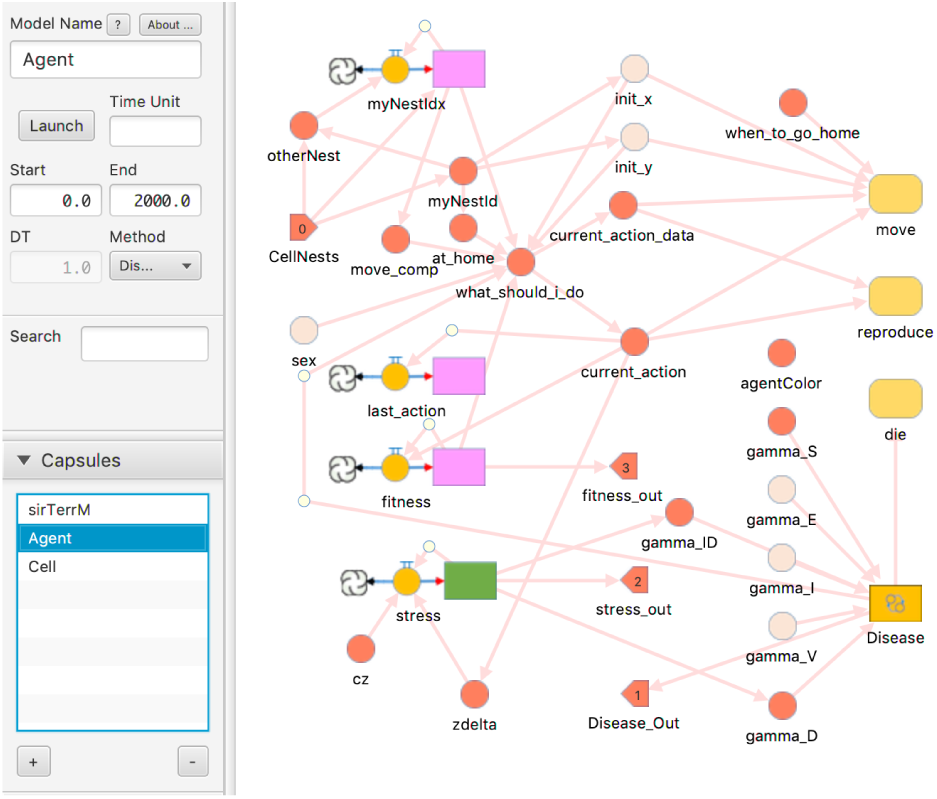

##### SimWorld

A *SimWorld* combines a CellMatrix with an AgentVector to create an artificial environment in which the Agents of the AgentVector range over the landscape created by the CellMatrix. Consequently, in addition to interacting with each other, Agents may determine the Cell in which they are located and interact with that or nearby Cells.

Pictured below is the top level Capsule of our model, `sirTerrM` showing the SimWorld labeled `NWorld`, which models a set of agents moving over a hexagonal cellular array. Other components in this Capsule are used to initialize the SimWorld and analyze and display results.

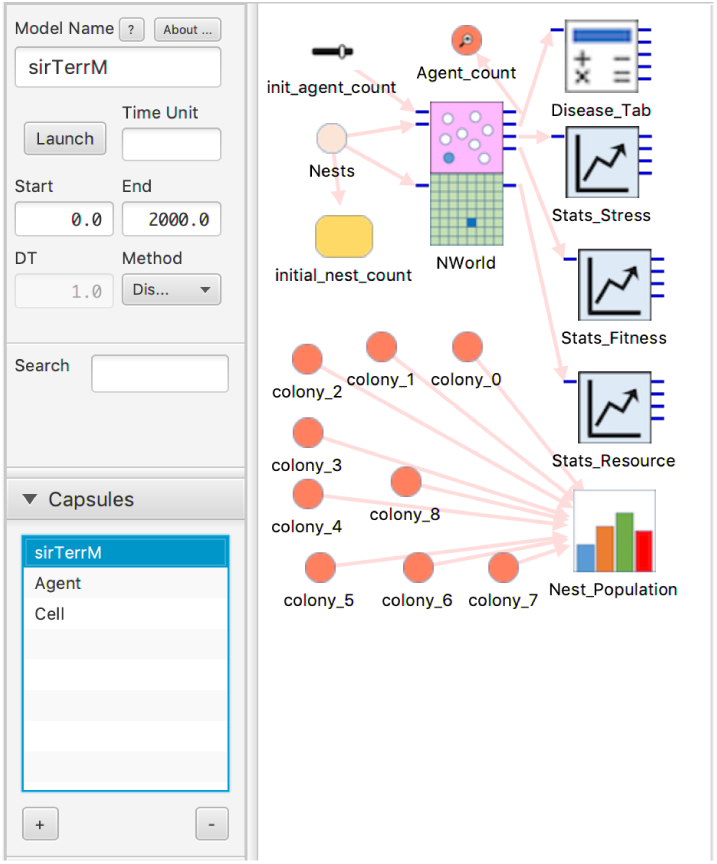

Numerus Model Builder also provides 2 additional Simulators for implementing a network spatial topology and using that topology as an agent landscape.

#### 2.2 Model Components

##### Stocks, Flows and Sequences

Traditional system dynamics modeling is centered on the stock-and-flow construct, whereby a stock, representing some dynamic model value, is updated by adding or subtracting the content of one or more flows (i.e respectively “flowing in or out” of the stock). In the simplest case, with a single flow flowing into a stock, the flow’s value is the derivative of the stock’s value over time. (When multiple flows are present, the derivative is found by summing the positive and negative flow values). The stock-flow model in system dynamics is especially apt for designing networks of stocks connected by flows that move values between the stocks.

Numerus fully supports the system dynamics paradigm, but adds several convenient modifications. A single Numerus Stock-Flow component may be substituted for a Stock with a single input Flow. This configuration can be particularly useful when modeling differential equations. Numerus also introduces a variation called Sequence, in which the Flow contains the Sequence’s *next* value, rather than content to be added to or subtracted from the current value. Sequences do not offer the Runge-Kutta integration algorithms available to Stocks to reduce the error when modeling continuous processes; therefore they are most useful for discrete models.

Numerus models are built using drag-and-drop gestures that place components on a design canvas. In the figures below, Stocks and Sequences respectively use lime-green and lavender-pink rectangles, and Flows are represented as yellow circles with arrows connecting to their Stocks.

The sirTerrM model uses both Stocks and Sequences in the Agent Capsule to represent continuous and discrete elements of the model.

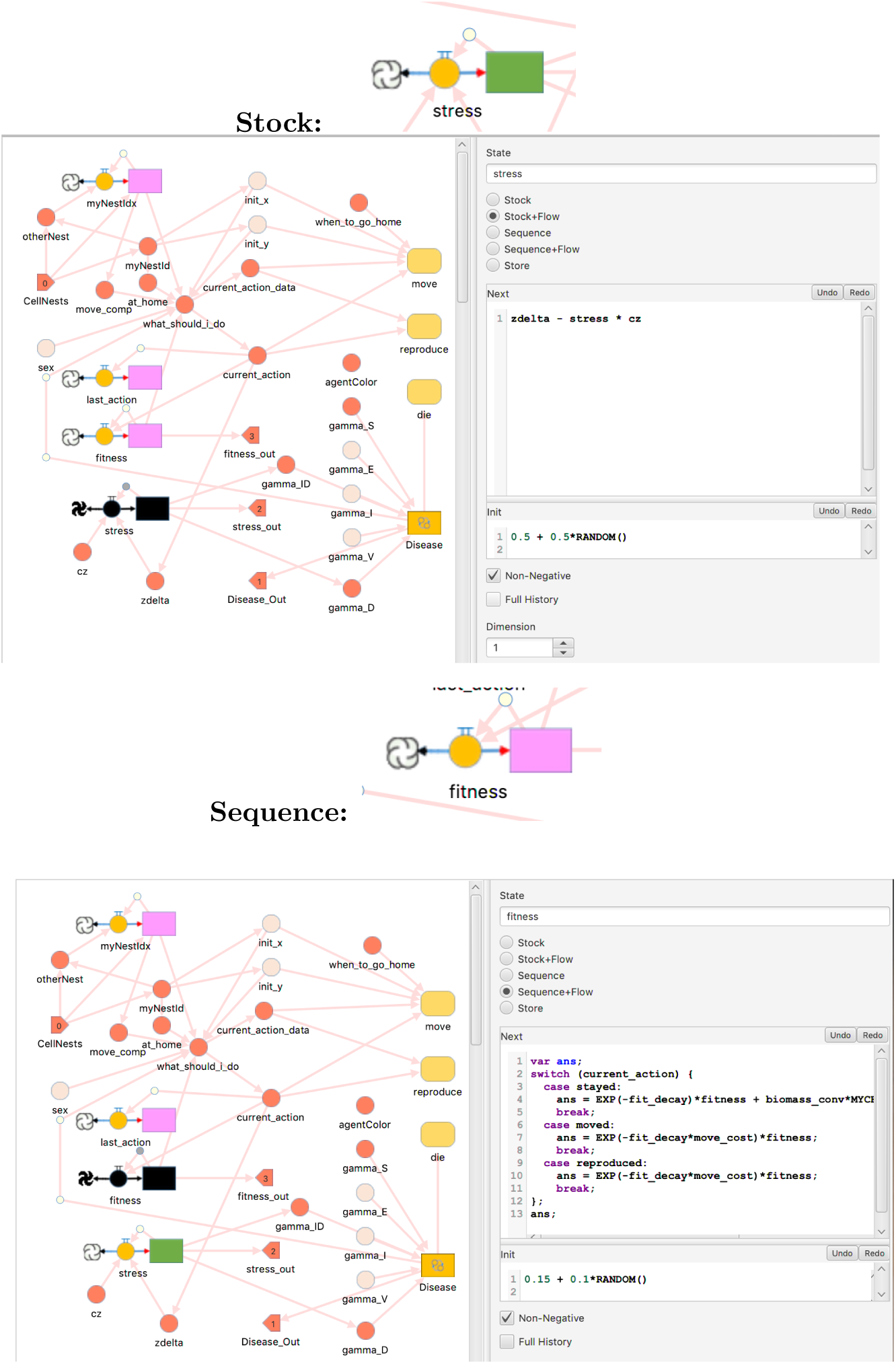

##### Discrete Value State

The *DiscreteState* component is similar to the Sequence, only its values are drawn from a small discrete set defined for use in the model. In the sirTerrM model, for example, the set of Disease states are represented by the letters S, E, I, V and D (for Susceptible, Exposed, Infected, Immune and Died, respectively). Other Discrete State sets identify agent sex (Male, Female), agent action (Stayed, Moved, Reproduced) and Cell type (Nest, Normal).

DiscreteState transitions are represented as a set of rules governing the transition from each allowable state. In this regard a DiscreteState component implements a finite automaton.

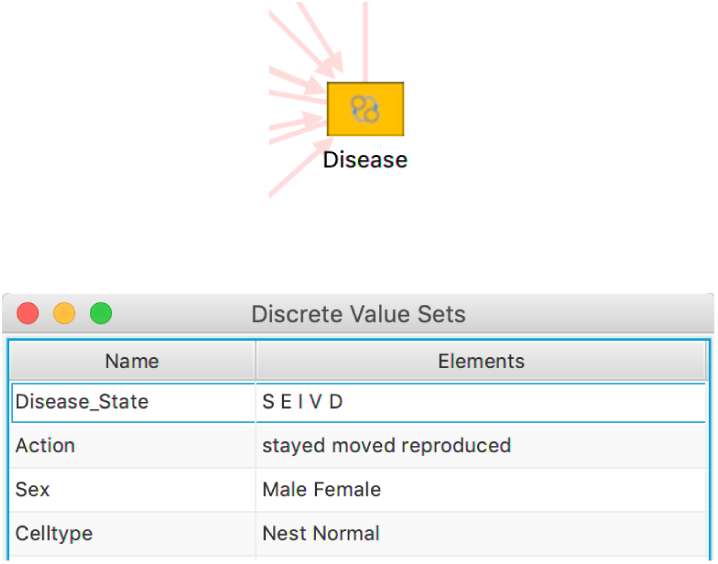

##### Graphical Display

Numerus provides a wide set of standard data display options (graphs, tables, etc.), including tableaux for visualizing the actions within cells and agents through the choice of color and (in the case of agents) size.

Certain analytical components also include graphs for visualizing their output. The Tabulator component determines at each point in time the number of agents or cells in a particular state. The Stats component computes statistical means and standard deviations for some state value across the set of cells or agents, and displays in a graph and table the running mean and mean ± standard deviation.

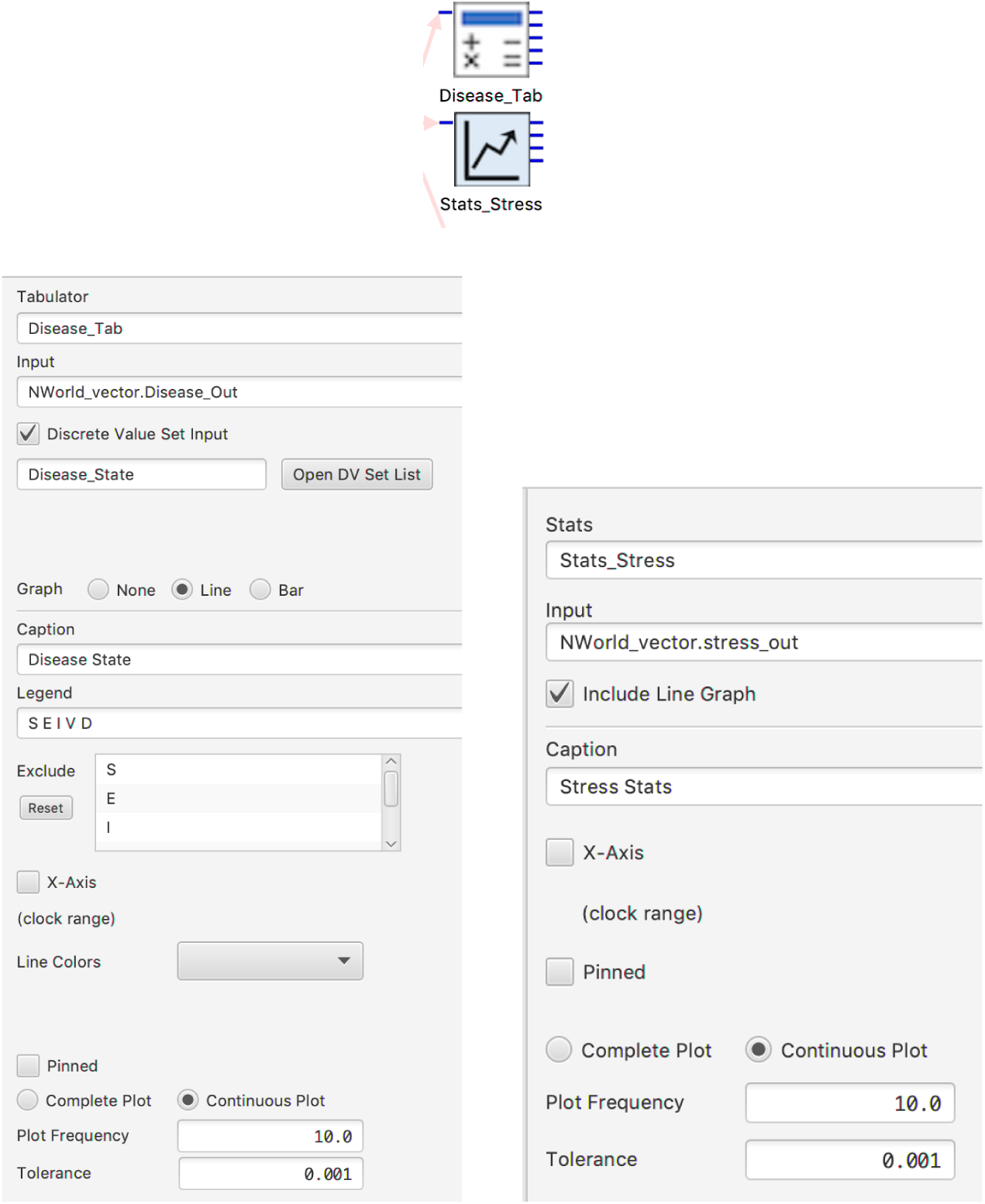

### 3 Building the Model

#### 3.1 Cell Layer

##### Cell Memory

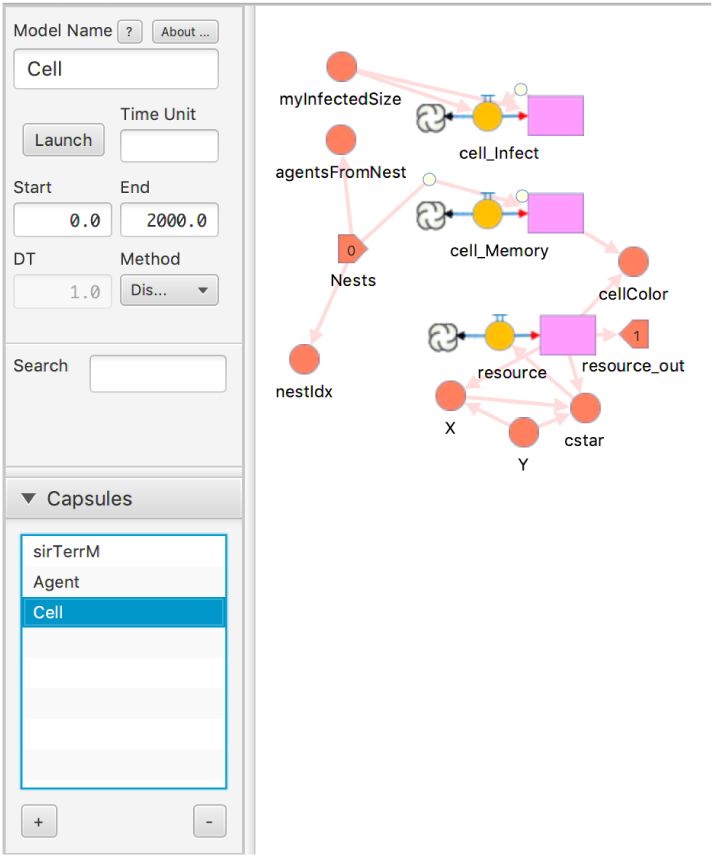

The population initially consists of *N*_0_ agents, but over time, due to births and deaths this number will change and be designated *N*_*t*_. To assign this, we go to Project >> Prelude.

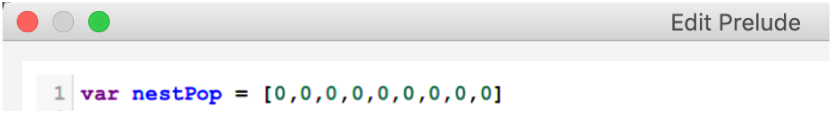

Recall that the quantities 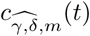, *m* = 1, …, *M* contain a fading memory of how many individuals from colony *m* have visited the cell 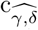 up until time *t*. In our implementation these data are stored in an *M* -element array in the Sequence `Cell Memory`. This array is initialized with each slot set to 0. At each point in time first the current value of each slot is aged by a factor of 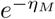 (where *η*_*M*_ is represented in the program by the parameter `mem fade`), and then the current population of the cell is distributed to the slots based on their nest of origin.

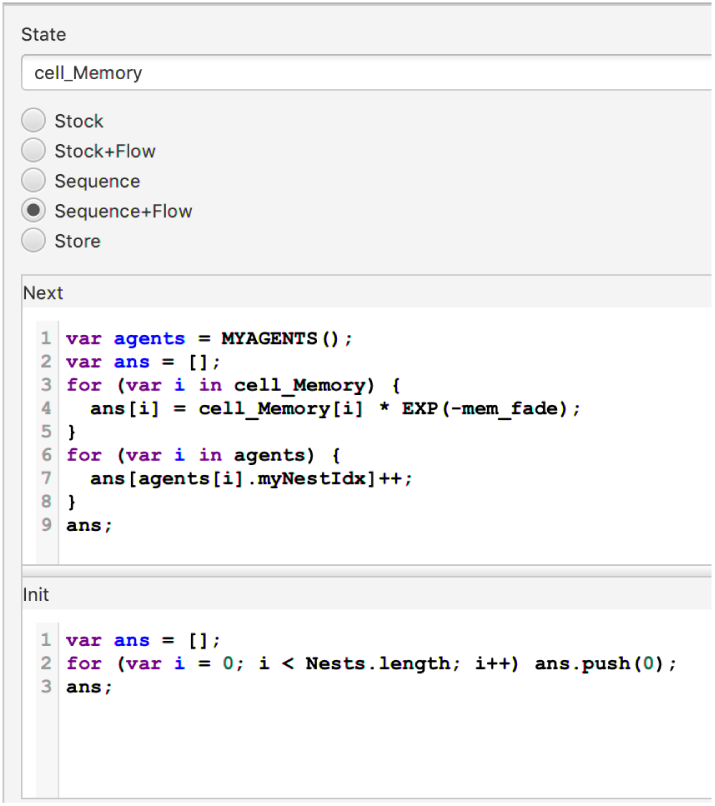

Note that `cell_Memory` is referenced by an agent during the process of determining its next move. This computation is contained in the agent’s `move_comp` term.

##### Cell Colors

We want the color of a cell in our visualization to reflect properties of the cell. Each nest is assigned a unique color which displays unless the nest is empty, in which case it displays in gray. Otherwise the cell’s color is computed to reflect the color of the nest of origin of the most number of visitors, or a neutral color if unvisited. This computation occurs in the Cell term `cellColor`.

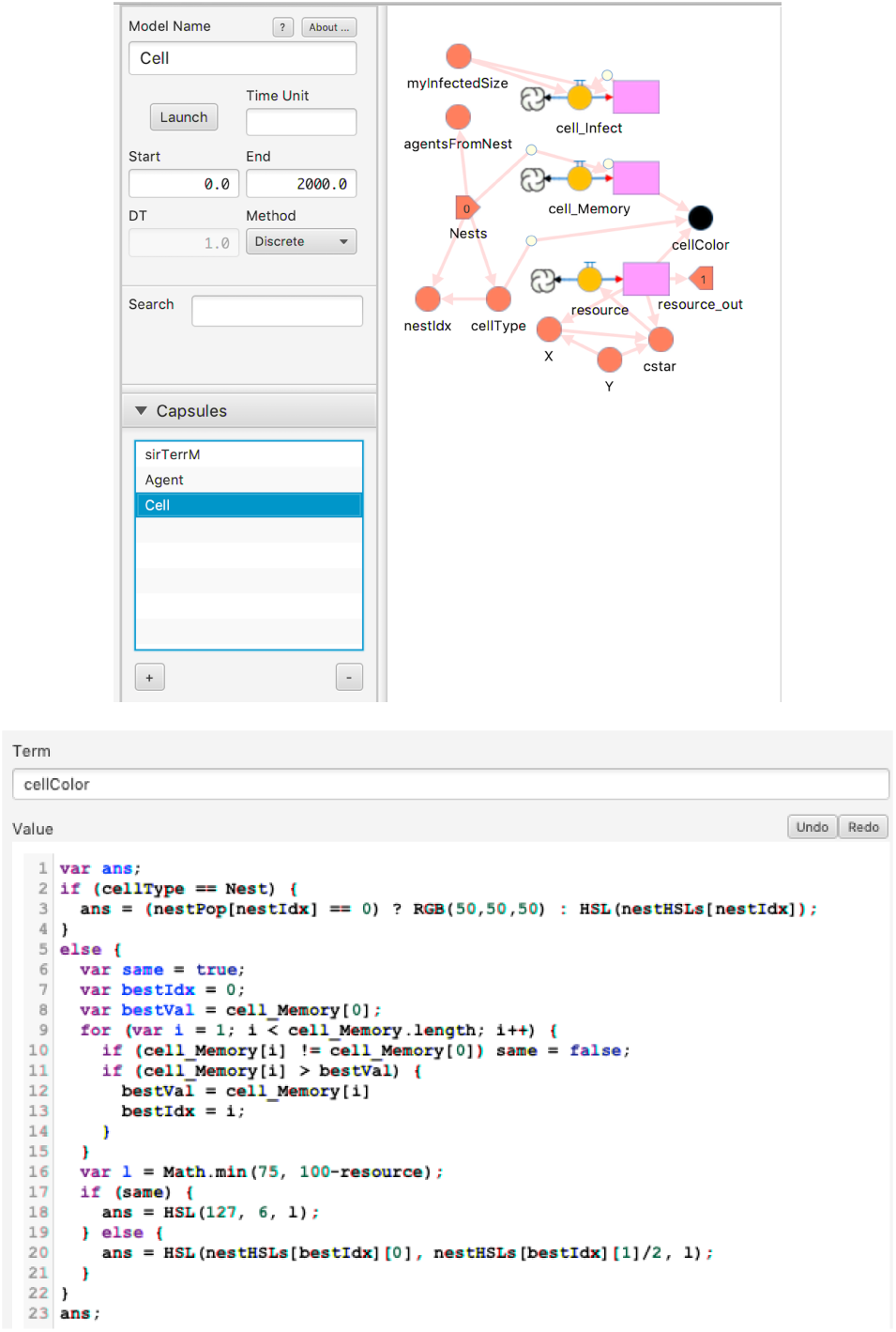

##### Disease Component

Underlying all dynamical systems modeling of epidemiological outbreaks and endemic diseases are formulations based on the concept of an SEIR progression. Susceptible individuals in disease class S enter disease class E on exposure to a pathogen (i.e., infected but not yet infectious themselves). After a period of latency, individuals in class E then transfer into the class of infectious individuals, I, only to transfer to a recovered or removed class R. However, note that once mortality is included in the model, then disease class R becomes ambiguous because removed individuals now include both recovered and dead individuals. Therefore, we use disease class V for recovered (i.e., “V” for naturally vaccinated individuals that have “recovered with immunity”) and D for dead. This SEIVD notation proves useful once SEIR processes are expanded to include births, recruitment, immigration, deaths, and emigration processes.

###### cell_Infect

Each cell has a `cell_Infect` Sequence representing the infectiousness of the cell at time *t* based on the number of infected individuals that have visited. We require a fade rate to account for an assumed decline in the infectiousness environment of the cell over time brought about by past visitors. If the fade rate is close to 0, this implies that an indirect transmission of pathogens via the cell’s environment is of significance, whereas if the fade rate goes towards ∞, then this implies that direct contact is increasingly important for transmission.

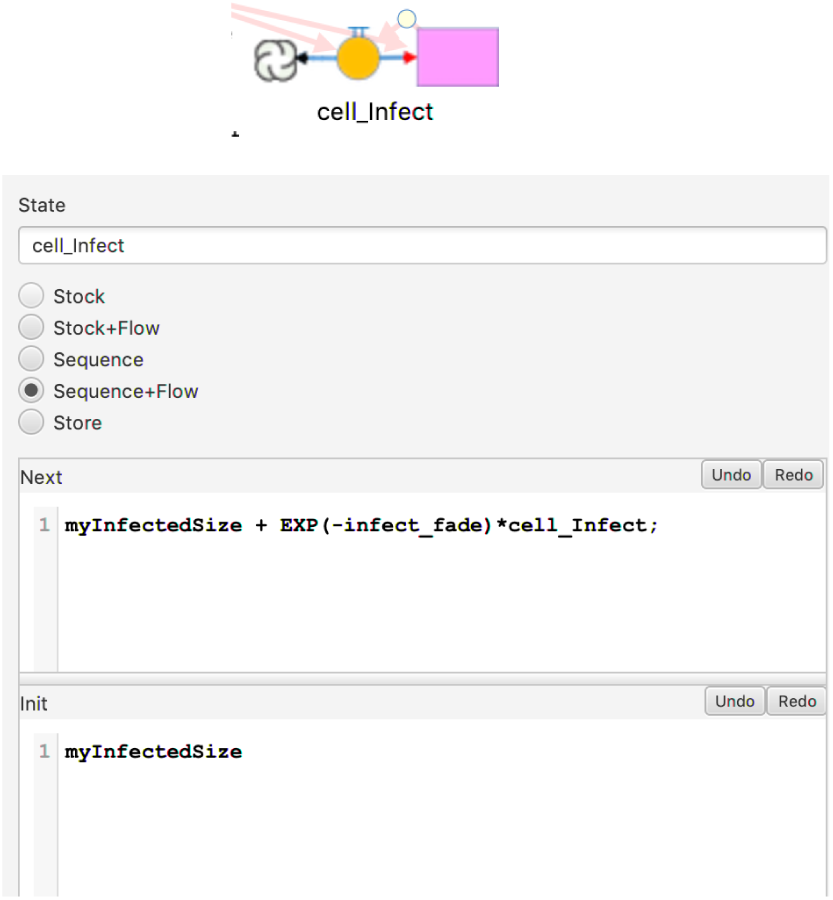

##### Resource

The third and final Cell state value is the resources it contains for nourishing the agents that visit. The formulae for determining resource dynamics involve both natural growth and agent consumption, and are discussed fully in the paper.

#### 3.2 Agent Layer

##### Disease Transition States

`Disease Transition Terms`: We include the 5 transition states as Terms: `gamma_`S, `gamma_`E, `gamma_`I, `gamma_`V, `gamma_`D, all of which govern the transitions of the Disease Value State (refer to Section 3 Model Components). Note that `gamma_`E, `gamma_`I, and `gamma_`V are constants derived from parameters. `gamma_`S, which governs the transition from Susceptible to Exposed, depends on the value of `cell_infect` (i.e. the infectiousness of the environment); and `gamma_`D, which governs the transition to Death, depends in part on agent stress.

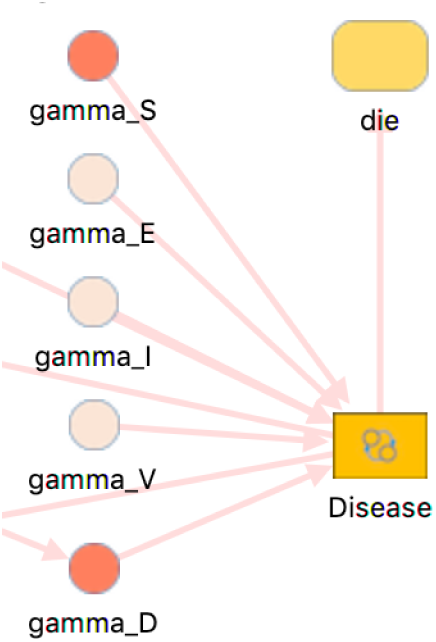

##### Movement

Determining movement is a computation that occurs in the Term `move_comp`. Section 6.2 of the manuscript explains movement in great detail. For the purpose of the Supplementary document, we will focus on a few fundamental concepts that determined our choosing of the terms and states for this particular component.

At the start of each time interval [*t, t* + 1], we initially make a case for whether or not individual *α* = 1, …, *N* remains in its current cell or moves to some other cell in the set `BLOCK`_*α*_(*r*), where *r* is some pre-selected positive integer that determines the furthest distance any single move can be.

We determined whether or not agent *α* belonging to colony *ℓ* would stay in its current cell or move to another one, based on either Distance Measure (`var dist`), Comparative Resource Measure (`var myResource`), and Comparative Occupation-History Measure (`var otherMaxOccupancy`).

There is also a stochastic trade-off with the agent movement rules, which play a role in determining cell attractiveness values.

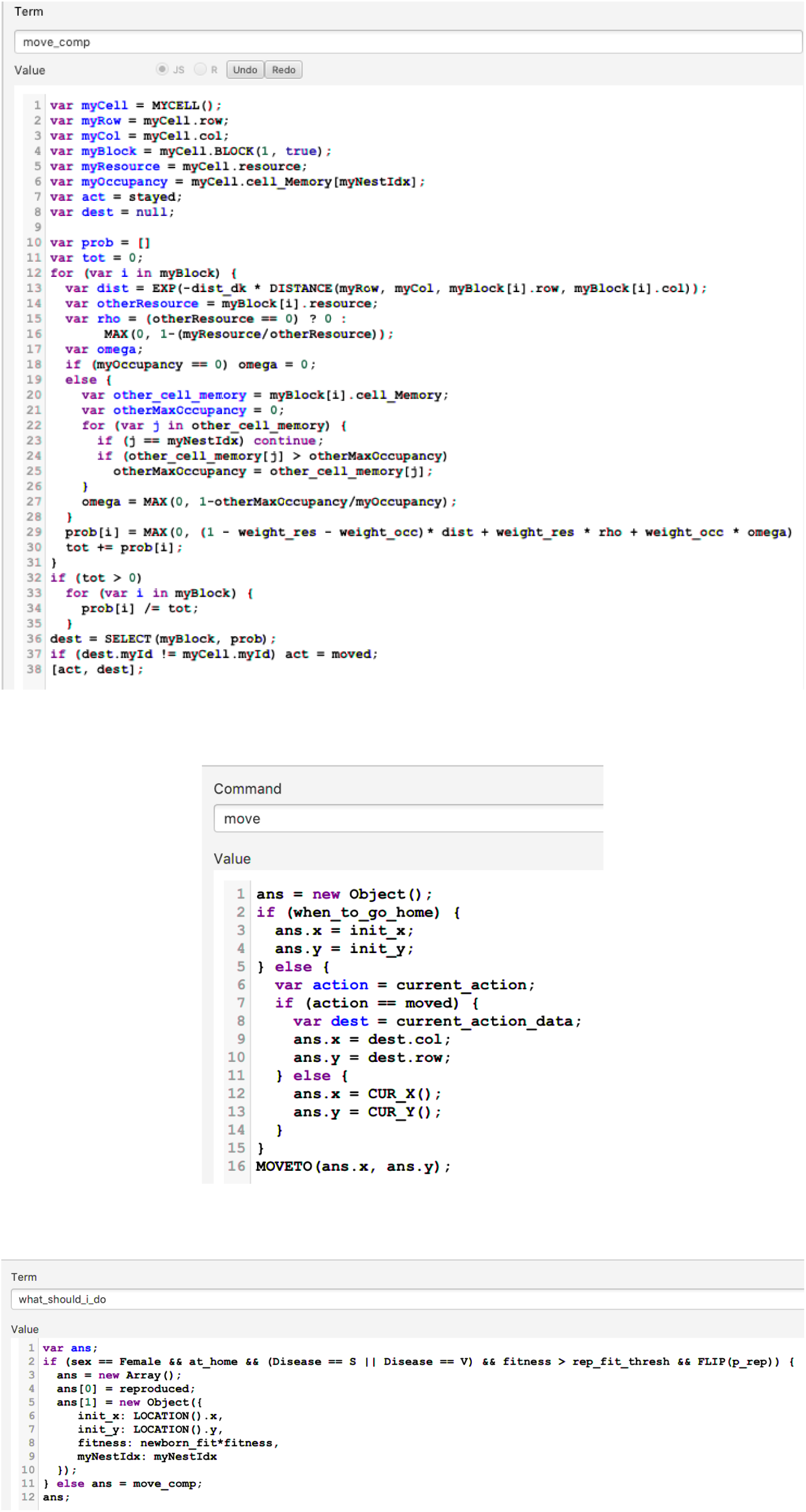

##### Agent Color

The Agent Color component also exists in order for us to observe distinctions between agents. It is determined by the agent Term `agentColor`, with each agent given the color associated with its home nest.

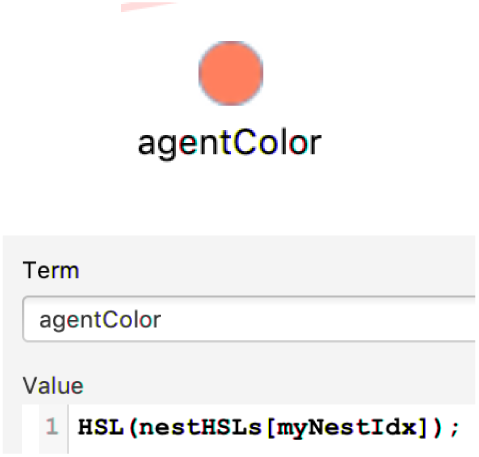

##### Fitness

A Fitness Sequence (defined in terms of the maximum resource extraction rate parameter) is also included in the model with the purpose of updating the stress state of individual agents. More specifically, the fitness of an individual is updated over the time interval[*t* − 1, *t*] for the maximum fitness value, the fitness decay factor, and the biomass conversion values, all of which are broken down thoroughly in Section 6.3, Ecological Dynamics, of the manuscript. If an individual moves out of a cell at time *t* − 1, then the fitness is reduced by a fraction that is some multiple of the decay rate.

We model Stress Dynamics through the agent’s Stress Stock, which contains the quantity *a*_*α*,5_(*t*) for *a* = 1, …, *N*, as described in Equation 11 of Section 6.3 in the paper. We note that the Stress component is initially given a random value between 0.5 and 1.

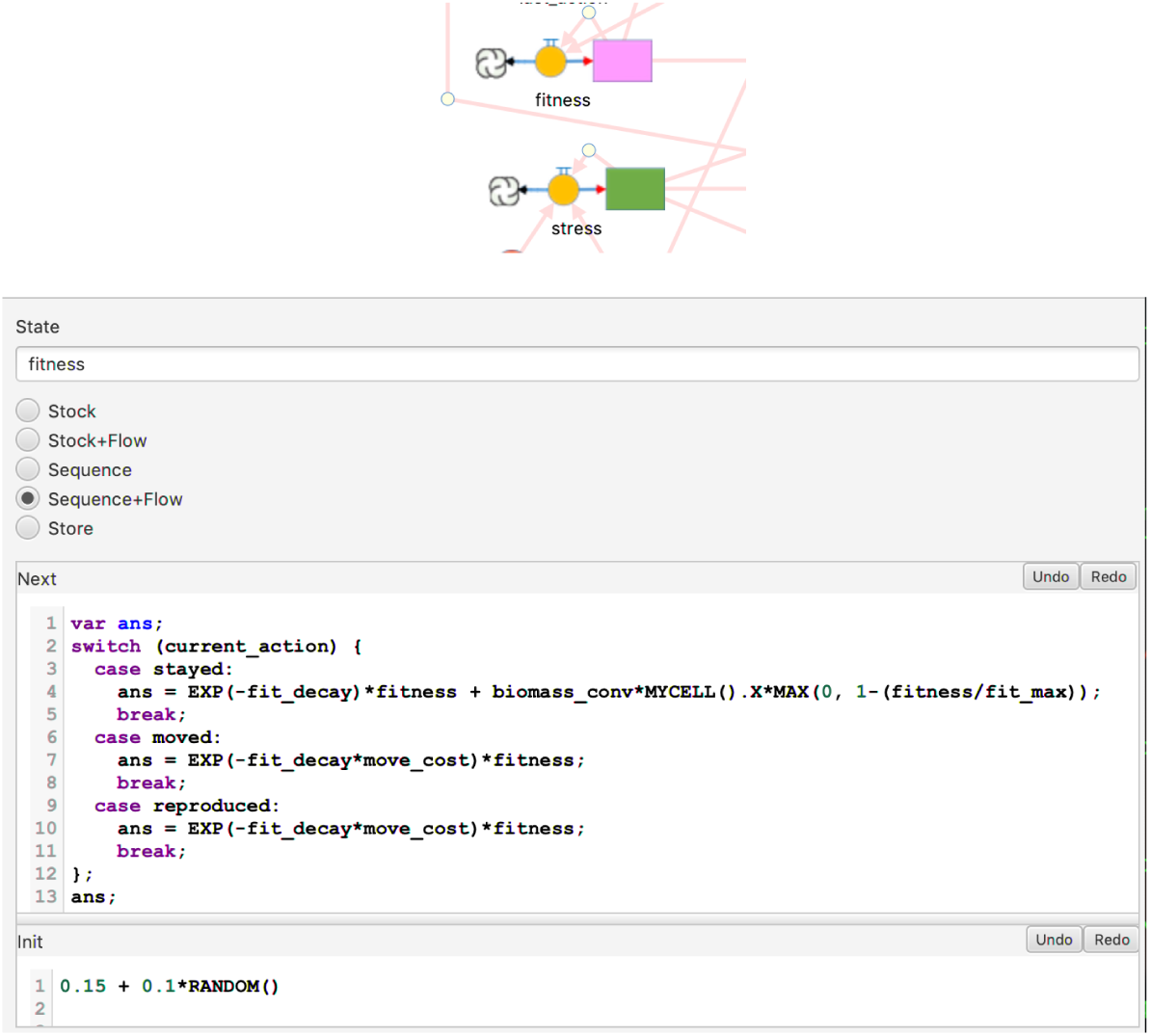

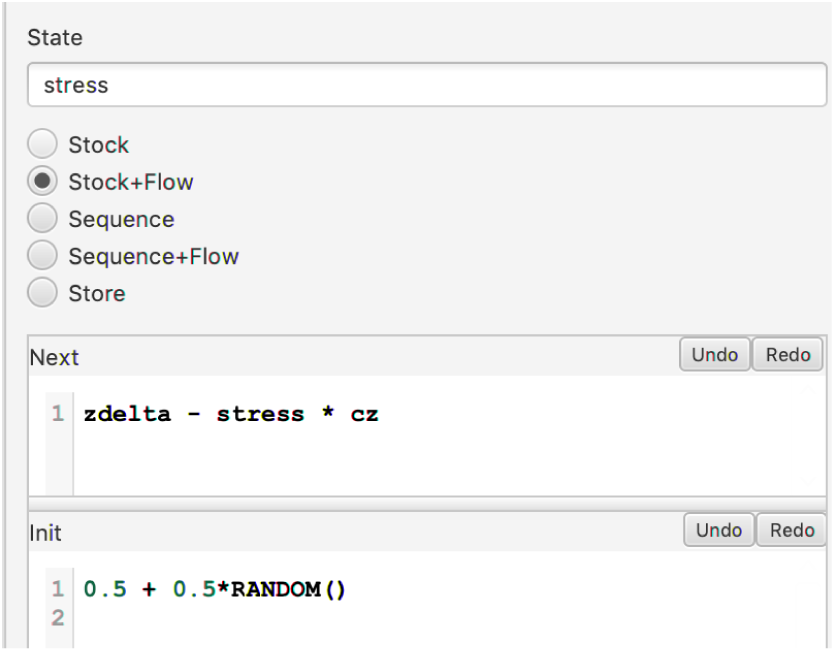

##### Reproduction

This model assumes nothing about mating structures and male limitations; therefore, only females reproduce, given a particular probability at regular colony-return times, provided they have sufficient fitness and sufficiently low stress values. Individuals die at natural-mortality specified rates, as well as with an additional disease-induced-mortality rate during their infectious period. Note that, as demonstrated in the figure below, Reproduction is a Command that occurs only when an agent selects “reproduced” as its current action. According to the rules of the model, this can occur only when the agent is in its home nest, which recurs on a cyclic basis (see the agent Term `what_should_i_do` for the rules governing this selection).

In our model, we incorporate the effect of pathogens on host population dynamics; therefore, individuals in disease states E and I do not reproduce. Individuals in disease state S or V, and whose fitness value exceeds *b*_min_ will not move at the next time step, but instead will give rise to a new individual. Furthermore, newly reproduced individuals will belong to the same colony as their parents.

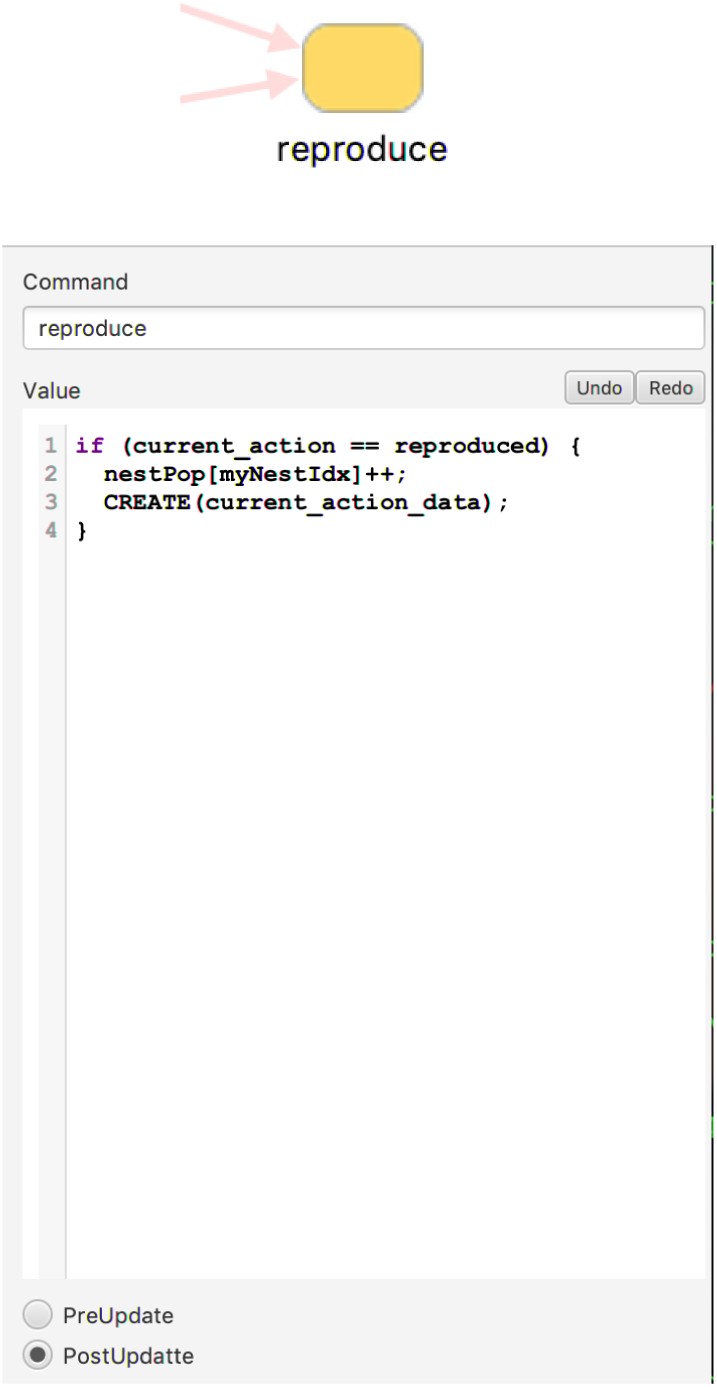

### 4 sirTerrM Layer

The top-most level includes the SimWorld container `NWorld` to house the cells and agents defined by the latter two Capsules.

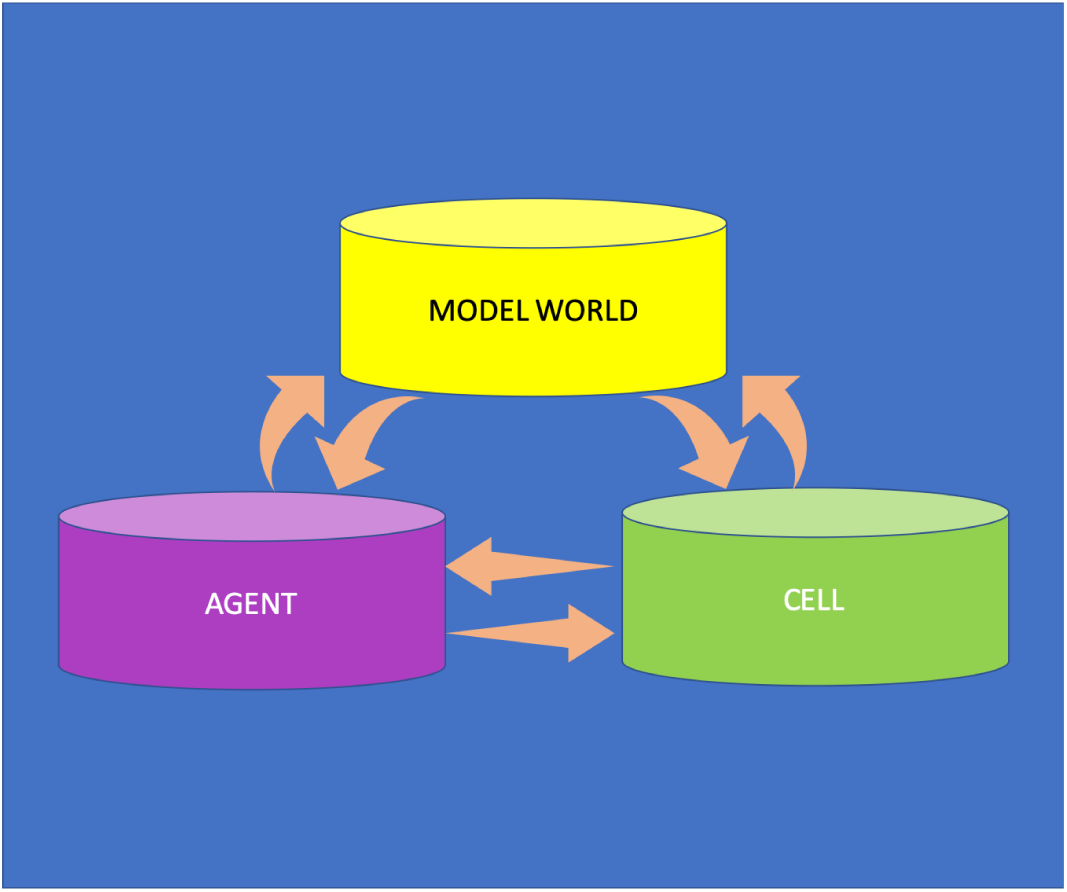

#### 4.1 Nests and Colonies

The Prelude contains the globally defined array `nestPop` that keeps track of the population of the 9 colonies used by the model. The Terms `colony_0` through `colony_8` each select a distinct slot of this array for inclusion in the bar graph `Nest_Population` (see below). The locations of the nests are defined in the Term `Nests` using the Numerus Cell ID notation C*Y* :*X*, where *Y* and *X* respectively indicate the row and column address of the cell. Nest locations are fed to the SimWorld through input pins to both its Agent and Cell constituents, since both need to be aware of these locations.

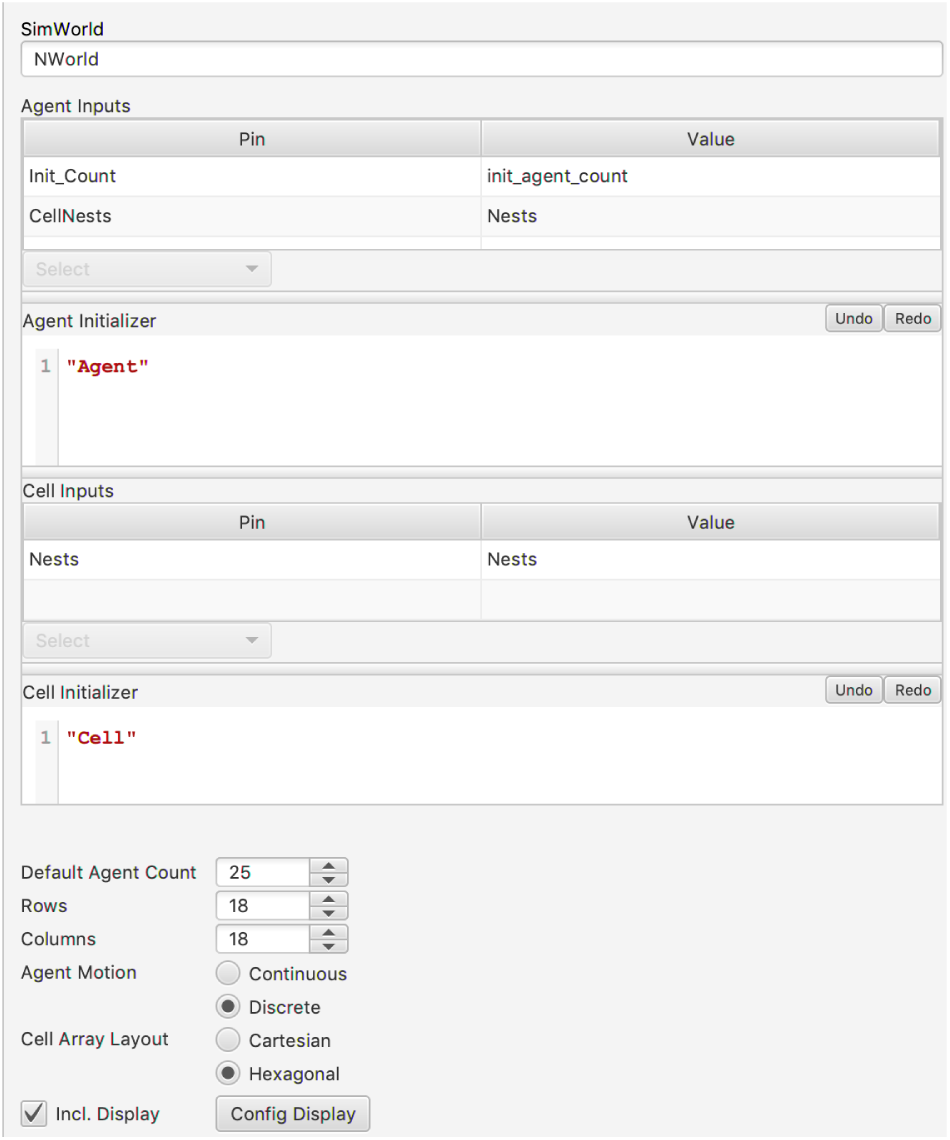

#### 4.2 Data Displays

There are a total of 6 displays showing the state of the running model. The first is a visualization of model activity generated by the `NWorld` SimWorld. The second and third are, respectively, the bar graph mentioned above showing the current colony populations, and line graphs tabulating the current disease state of the agents. The final three displays provide statistical information across the agent and cell populations for Stress and Fitness (agent); and Resource (cell). The Stats plug-in is used to create graphs showing the mean and mean ± standard deviation of each of these quantities.

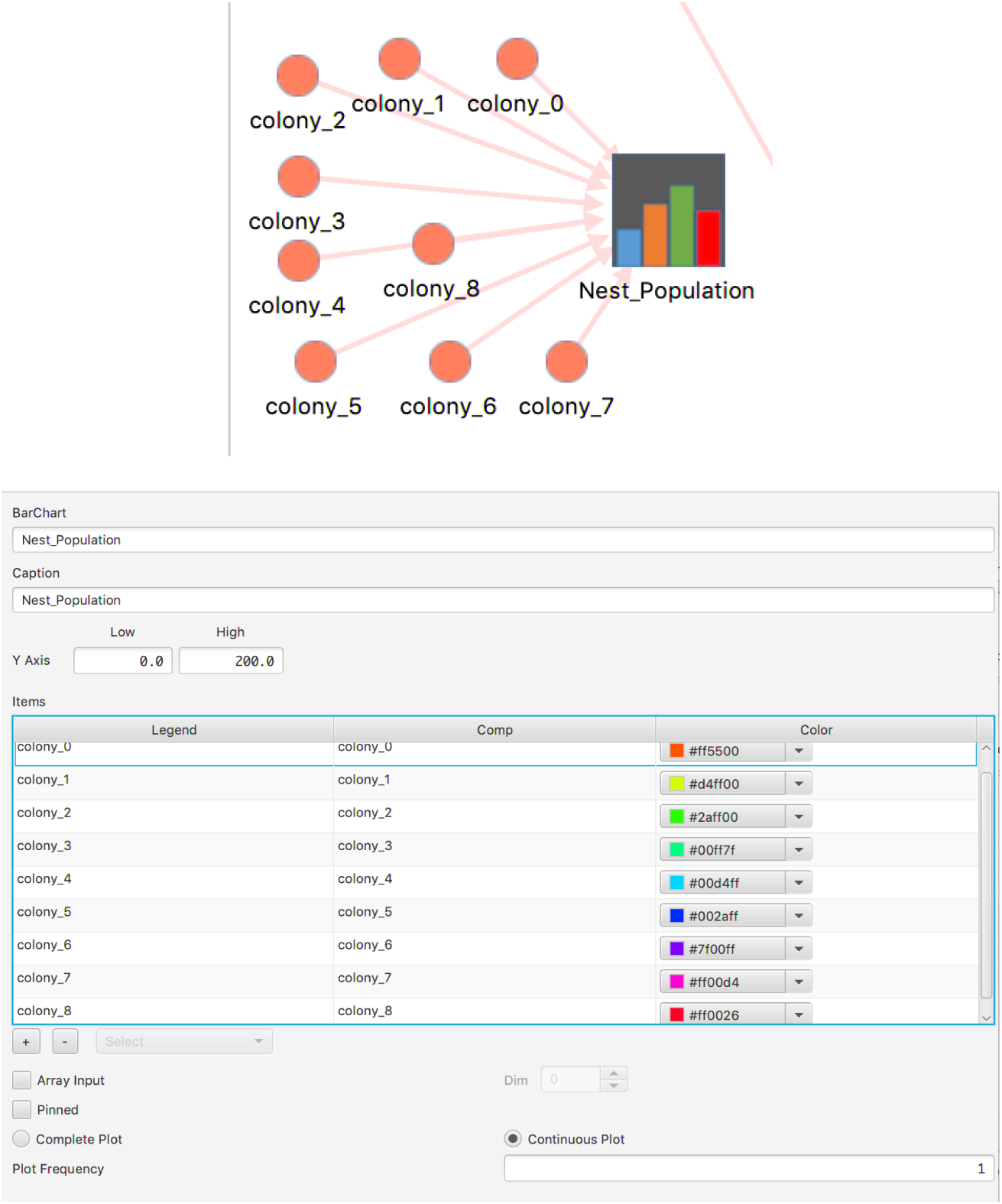

### 5 Text-to-Model Terminology Translation

The tables below provide a translation between the notation used in the text and that employed in the model.

**Table 1:**
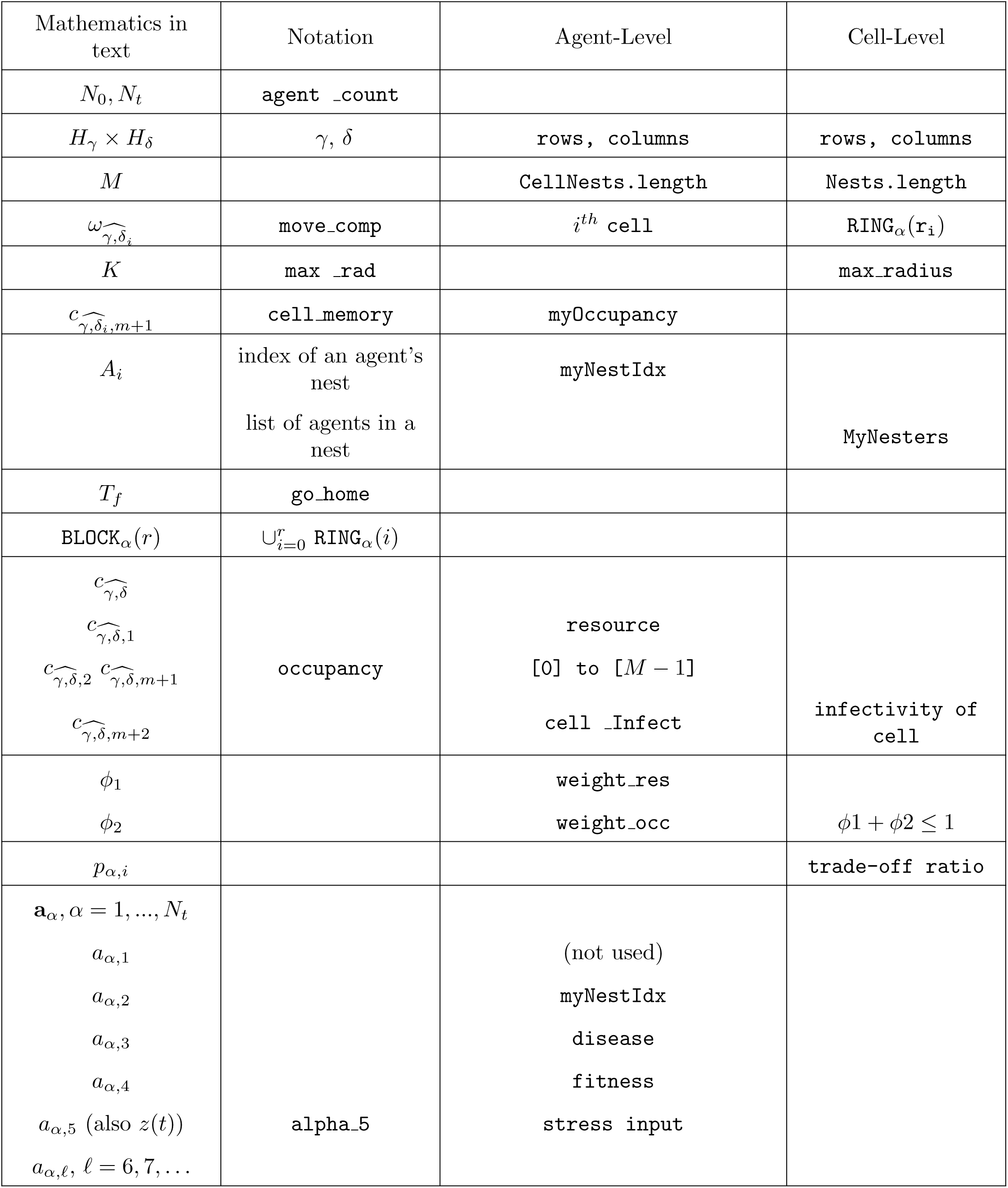
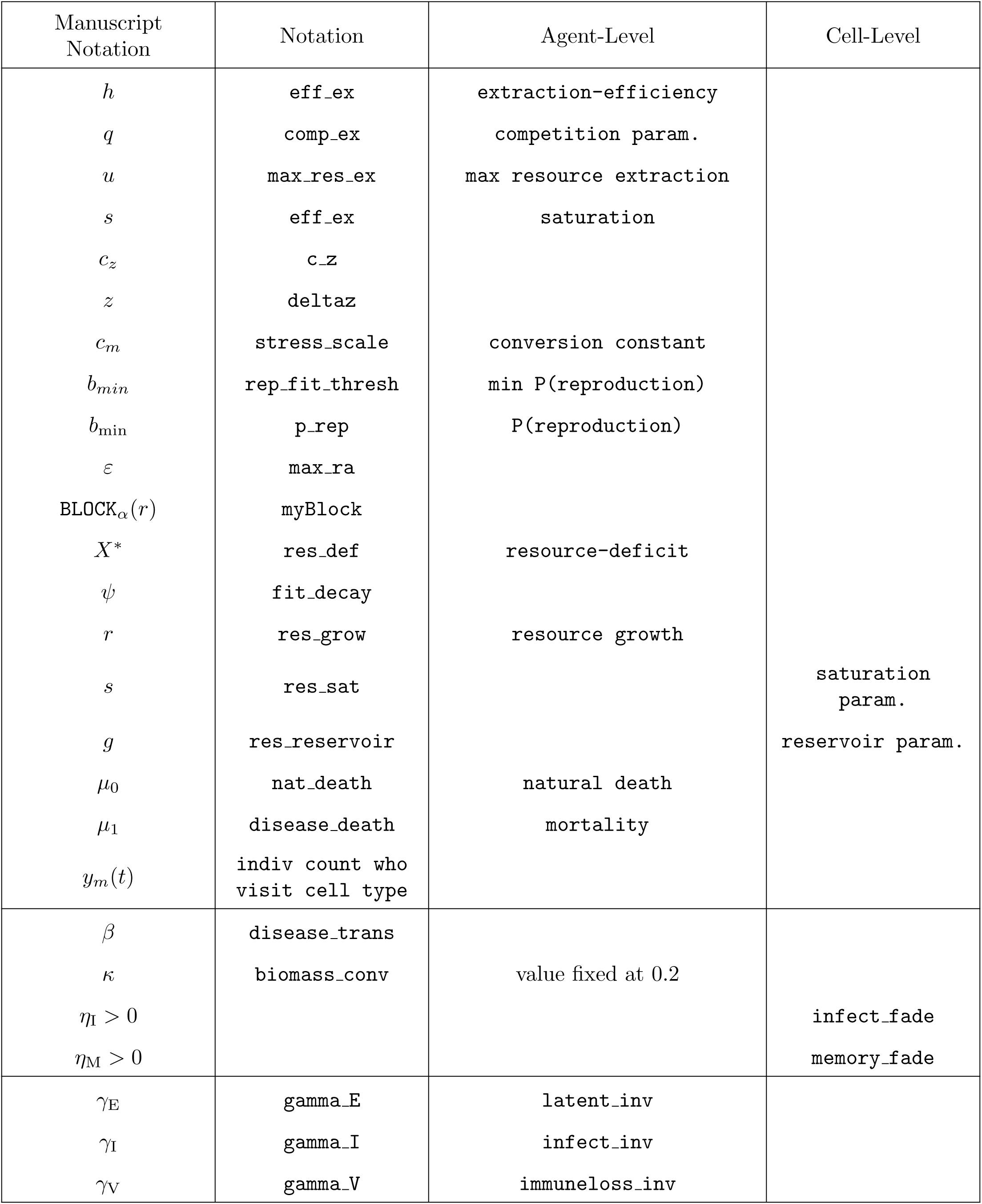
Translation of notation between mathematics in text and NMB code

### 6 NovaScript

The NovaScript generated by NMB is attached to the end of this document. The model file is called GetzEtAlPLoSOct2019.nmd, which can be downloaded at https://www.dropbox.com/s/24j6js6js6h5ur3/GetzEtAlPLoSOct2019.nmd?dl=0 and can be run after downloading a version of NMB at numerusinc.com

~~~
var fit_max = 1;
var rho_crit = ø.3;
var biomass_conv = ø.2;
var cycle_home = 10;
var eff_ex = 5;
var max_res_ex = 5;
var res_sat = 20;
var res_reservoir = ø.1;
var stress_scale = ø.5;
var prob_f = ø.6;
var p_rep = ø.8;
var parent_fit = ø.65;
var newborn_fit = ø.3;
var max_rad = 2;
var mem_fade = ø.3;
var dist_dk = ø.3;
var infect_fade = ø.5;
var latent_inv = ø.5;
var infect_inv = ø.2;
var immunelose_inv = ø.02;

defineSchema(“Cell”, {
    specifies: “Capsule”,
    methods: {
    },
    components: {
       cell_Memory : {
         specifies: “Sequence”,
         count: 1,
         nonNegative: false,
         history: 100,
         initial: function(){
              var ans = [];
              for (var i = 0; i < this.Nests.length; i++)
ans.push(0);
              return ans;
              },
         next: function(){
              var agents = this.MYAGENTS();
              var ans = [];
              for (var i in this.cell_Memory) {
              ans[i] = this.cell_Memory[i] * EXP(-this.mem_fade);
              }
              for (var i in agents) {
              ans[agents[i].myNestIdx]++;
              }
              return ans;
              },
     },
     resource : {
              specifies: “Sequence”,
              count: 1,
              nonNegative: false,
              history: 100,
              initial: ‘2+this.RANDOM(1)’,
              next: ‘this.cstar + this.res_grow * (1 - this.cstar/ this.res_sat)*(this.cstar + this.res_reservoir)’,
     },
              cell_Infect : {
              specifies: “Sequence”,
              count: 1,
              nonNegative: false,
              history: 100,
              initial: ‘this.myInfectedSize’,
              next: ‘this.myInfectedSize + EXP(-
this.infect_fade)*this.cell_Infect;’,
              },
              cellType : {
              specifies: “Term”,
              exp: ‘(this.Nests.indexOf(this.myId) >= 0) ? Nest :
Normal’,
              },
              nestIdx : {
              specifies: “Term”,
              exp: ‘(this.cellType == Nest) ?
this.Nests.indexOf(this.myId) : -1;’,
              },
              Y : {
              specifies: “Term”,
              exp: function(){
              var agts = this.MYAGENTS();
              agts = _.filter(agts, function(x){return x.last_action
== stayed;});
              var count = agts.length;
              return count;
              },
           },
           cstar : {
              specifies: “Term”,
              exp: ‘this.resource - this.Y*this.X’,
           },
           cellColor : {
              specifies: “Term”,
              exp: function(){
                 var ans;
                 if (this.cellType == Nest) {
                  ans = (nestPop[this.nestIdx] == 0) ? RGB(50,50,50) :
HSL(nestHSLs[this.nestIdx]);
              }
              else {
              var same = true;
              var bestIdx = 0;
              var bestVal = this.cell_Memory[0];
              for (var i = 1; i < this.cell_Memory.length; i++) {
              if (this.cell_Memory[i] != this.cell_Memory[0])
same = false;
              if (this.cell_Memory[i] > bestVal) {
                bestVal = this.cell_Memory[i]
                bestIdx = i;
                }
              }
              if (same) {
                var l = 50-(this.resource/2);
                ans = HSL(127, 85, l);
              } else {
                var l = Math.min(75, 100-this.resource);
                ans = HSL(nestHSLs[bestIdx][0], nestHSLs[bestIdx]
[1]/2, l);
                }
              }
              return ans;
              },
          },
          myInfectedSize : {
              specifies: “Term”,
              exp: ‘this.COUNT(function(agt){return agt.Disease == I ||
agt.Disease == E;}, this.MYAGENTS());’,
              },
              Nests : {
                   specifies: “InPin”,
                   exp: 0.0,
              },
              agentsFromNest : {
                   specifies: “Term”,
                   exp: function(){
              var ans = []
              for (var i = 0; i < this.Nests.length; i++)
ans.push(0);
              var agents = this.MYAGENTS();
              for (var idx in agents) {
                 agent = agents[idx];
                 var nestIdx = agent.myNestIdx;
                 ans[nestIdx]++;
              }
              return ans;
              },
           },
              resource_out : {
                  specifies: “OutPin”,
              exp: ‘this.resource’,
     },
    X : {
              specifies: “Term”,
              exp: function(){
                  var ans = ø.0;
                  if (this.Y > 0) {
                   var v0 = this.resource/this.Y;
                   var v1 = (this.max_res_ex * this.resource)/
(this.eff_ex * (1 + this.comp_ex * (this.Y-1)) + this.resource);
                 ans = MIN(v0, v1);
              }
              return ans;
              },
              

      },
              disease_trans : {
                 specifies: “InPin”,
                 exp: “mscope.disease_trans”,
              },
              disease_death : {
                 specifies: “InPin”,
                 exp: “mscope.disease_death”,
              },
              p_switch : {
                 specifies: “InPin”,
                 exp: “mscope.p_switch”,
              },
              nat_death : {
                 specifies: “InPin”,
                 exp: “mscope.nat_death”,
              },
              weight_res : {
                 specifies: “InPin”,
                 exp: “mscope.weight_res”,
              },
              weight_occ : {
                 specifies: “InPin”,
                 exp: “mscope.weight_occ”,
              },
              res_grow : {
                 specifies: “InPin”,
                 exp: “mscope.res_grow”,
              },
              comp_ex : {
                 specifies: “InPin”,
                 exp: “mscope.comp_ex”,
              },
              rep_fit_thresh : {
                 specifies: “InPin”,
                 exp: “mscope.rep_fit_thresh”,
              },
              res_def : {
                 specifies: “InPin”,
                 exp: “mscope.res_def”,
              },
              stress_decay : {
                 specifies: “InPin”,
                 exp: “mscope.stress_decay”,
              },
              stress_death : {
                 specifies: “InPin”,
                 exp: “mscope.stress_death”,
              },
              fit_decay : {
                 specifies: “InPin”,
                 exp: “mscope.fit_decay”,
              },
              move_cost : {
                 specifies: “InPin”,
                 exp: “mscope.move_cost”,
              },
              },
              });
defineSchema(“Agent”, {
              specifies: “Capsule”,
              methods: {
              },
              components: {
                  fitness : {
                     specifies: “Sequence”,
                     count: 1,
                     nonNegative: true,
                     history: 100,
                     initial: ‘0.15 + 0.1*this.RANDOM()’,
                     next: function(){
                        var ans;
                        switch (this.current_action) {
                        case stayed:
                         ans = EXP(-this.fit_decay)*this.fitness +
this.biomass_conv*this.MYCELL().X*MAX(0, 1-(this.fitness/
this.fit_max));
                        break;
                     case moved:
                        ans = EXP(-
this.fit_decay*this.move_cost)*this.fitness;
              break;
              case reproduced:
              ans = EXP(-
this.fit_decay*this.move_cost)*this.fitness;
                  break;
              };
              return ans;
              },
         },
         stress : {
              specifies: “Variable”,
              count: 1,
              nonNegative: true,
              history: 100,
              initial: ‘0.5 + 0.5*this.RANDOM()’,
              prime: ‘this.zdelta - this.stress * this.cz’,
          },
              last_action : {
                 specifies: “Sequence”,
              count: 1,
              nonNegative: false,
              history: 100,
              initial: ‘stayed’,
              next: ‘this.current_action’,
           },
              myNestIdx : {
                 specifies: “Sequence”,
              count: 1,
              nonNegative: false,
              history: 100,
              initial: ‘FLOOR(this.RANDOM(this.CellNests.length))’,
              next: function(){
                  var ans = this.myNestIdx;
                  if (this.p_switch > 0) {
                   if (this.otherNest >= 0 &&
                     nestPop[this.otherNest]/nestPop[this.myNestIdx]
< this.rho_crit &&
                 this.FLIP(this.p_switch)) {
              ans = this.otherNest;
              nestPop[this.myNestIdx]--;
              nestPop[this.otherNest]++;
              };
            }
            return ans
            },
         },
              gamma_ID : {
                 specifies: “Term”,
                 exp: ‘(this.nat_death + this.disease_death) * (1 +
this.stress_death*this.stress);’,
              },
              gamma_D : {
                 specifies: “Term”,
                 exp: ‘this.nat_death * (1 +
this.stress_death*this.stress);’,
              },
              gamma_V : {
                 specifies: “Term”,
              constant: true,
                 exp: ‘this.immunelose_inv’,
              },
              gamma_I : {
                 specifies: “Term”,
              constant: true,
                 exp: ‘this.infect_inv’,
              },
              gamma_E : {
                 specifies: “Term”,
              constant: true,
                 exp: ‘this.latent_inv’,
              },
              gamma_S : {
                 specifies: “Term”,
                 exp: ‘this.disease_trans * this.MYCELL().cell_Infect’,
              },
              when_to_go_home : {
                 specifies: “Term”,
                 exp: ‘this.TIME()%this.cycle_home == 0’,
              },
              myNestId : {
                 specifies: “Term”,
                 exp: ‘this.CellNests[this.myNestIdx]’,
              },
              agentColor : { specifies: “Term”,
                 exp: ‘HSL(nestHSLs[this.myNestIdx]);’,
              },
              init_x : {
                 specifies: “Term”,
              constant: true,
                 exp: ‘this.COL_OF(this.myNestId)’,
              },
              init_y : {
                 specifies: “Term”,
              constant: true,
                 exp: ‘this.ROW_OF(this.myNestId)’,
              },
              CellNests : {
                 specifies: “InPin”,
              exp: 0.0,
              },
              Disease_Out : {
                 specifies: “OutPin”,
                 exp: ‘this.Disease’,
              },
              cz : {
                 specifies: “Term”,
                 exp: ‘this.stress_decay’,
              },
              zdelta : {
                 specifies: “Term”,
              exp: function(){
              var ans;
              switch (this.current_action) {
              case stayed:
              case reproduced:
              ans = MAX(0, this.stress_scale * (this.res_def -
this.MYCELL().X));
                    break;
                  case moved:
                    ans = this.stress_scale * this.res_def;
                    break;
              };
              return ans;
              },
          },
          at_home : {
                 specifies: “Term”,
                 exp: ‘this.myNestId == this.MYCELL().myId’,
          },
           what_should_i_do : {
                 specifies: “Term”,
              exp: function(){
                   var ans;
                   if (this.sex == Female && this.at_home &&
(this.Disease == S || this.Disease == V) && this.fitness >
this.rep_fit_thresh && this.FLIP(this.p_rep)) {
              ans = new Array();
              ans[0] = reproduced;
              ans[1] = new Object({
                  init_x: this.LOCATION().x,
                  init_y: this.LOCATION().y,
                 fitness: this.newborn_fit*this.fitness,
                 myNestIdx: this.myNestIdx
              });
             } else ans = this.move_comp;
             return ans;
             },
          },
          current_action : {
                 specifies: “Term”,
                 exp: ‘this.what_should_i_do[0]’,
          },
           current_action_data : {
                 specifies: “Term”,
                 exp: ‘this.what_should_i_do[1]’,
              },
              stress_out : {
                 specifies: “OutPin”,
                 exp: ‘this.stress’,
              },
              fitness_out : {
                 specifies: “OutPin”,
                 exp: ‘this.fitness’,
              },
              move_comp : {
                 specifies: “Term”,
                 exp: function(){
                   var myCell = this.MYCELL();
                   var myRow = myCell.row;
                   var myCol = myCell.col;
                   var myBlock = myCell.BLOCK(1, true);
                   var myResource = myCell.resource;
                   var myOccupancy = myCell.cell_Memory[this.myNestIdx];
                   var act = stayed;
                   var dest = null;
              

                   // rule 1
                   var prob = []
                   var tot = 0;
                   for (var i in myBlock) {
                      var dist = EXP(-this.dist_dk * this.DISTANCE(myRow,
myCol, myBlock[i].row, myBlock[i].col));
              var otherResource = myBlock[i].resource;
              var rho = (otherResource == 0) ? 0 :
                 MAX(0, 1-(myResource/otherResource));
              var omega;
              if (myOccupancy == 0) omega = 0;
              else {
                  var other_cell_memory = myBlock[i].cell_Memory;
                  var otherMaxOccupancy = 0;
                  for (var j in other_cell_memory) {
                   if (j == this.myNestIdx) continue;
                   if (other_cell_memory[j] > otherMaxOccupancy)
                     otherMaxOccupancy = other_cell_memory[j];
              }
                 omega = MAX(0, 1-otherMaxOccupancy/myOccupancy);
              }
              //prob[i] = MAX(0, (1 - (weight_res+weight_occ) *
dist + weight_res * rho + weight_occ * omega))
              prob[i] = MAX(0, (1 - this.weight_res –
this.weight_occ)* dist + this.weight_res * rho + this.weight_occ *
              omega)
              tot += prob[i];
              }
              if (tot > 0)
              for (var i in myBlock) {
              prob[i] /= tot;
                 }
              dest = this.SELECT(myBlock, prob);
              if (dest != myCell) act = moved;
              return [act, dest];
              },
          },
          sex : {
                 specifies: “Term”,
              constant: true,
                 exp: ‘(this.FLIP(this.prob_f)) ? Female : Male’,
           },
          otherNest : {
                 specifies: “Term”,
                 exp: function(){
                  found = -1;
                  var myCell = this.MYCELL().myId;
                  if (myCell != this.myNestId) {
                   for (var i in this.CellNests) {
                    if (this.CellNests[i] == myCell) {
                     found = i;
                     break;
                  }
                }
              };
              return found;
              },
    },
    move : {
             specifies: “Command”,
             when: “post”,
             exp: function(){
                 ans = new Object();
                 if (this.when_to_go_home) {
                   ans.x = this.init_x;
                   ans.y = this.init_y;
              } else {
                var action = this.current_action;
                if (action == moved) {
                 var dest = this.current_action_data;
                   ans.x = dest.col;
                   ans.y = dest.row;
              } else {
                 ans.x = this.CUR_X();
                 ans.y = this.CUR_Y();
                }
              }
         this.MOVETO(ans.x, ans.y);
              }
         },
         reproduce : {
                 specifies: “Command”,
              when: “post”,
              exp: function(){
                  if (this.current_action == reproduced) {
                  nestPop[this.myNestIdx]++;
                  this.CREATE(this.current_action_data);
              }
              }
         },
         die : {
              specifies: “Command”,
              when: “post”,
              exp: function(){
                if (this.Disease == D) {
                  this.Disease == S;
                  nestPop[this.myNestIdx]--;
                  this.KILL(this.myId);
              }
              }
           },
           Disease : {
              specifies: “DiscreteState”,
              dsGroup: “Disease_State”,
              stateSize: 5,
              stateNames: [“S”, “E”, “I”, “V”, “D”],
              initial: ‘(this.myId < 1) ? E : S’,
              next: {
              S: ‘this.SELECT([E,D,S],\n’+
              ‘[this.gamma_S*(1-EXP(-this.gamma_S –
this.gamma_D))/(this.gamma_S + this.gamma_D),\n’+
              ‘this.gamma_D*(1-EXP(-this.gamma_S –
this.gamma_D))/(this.gamma_S + this.gamma_D),\n’+
              ‘EXP(-this.gamma_S - this.gamma_D)]);\n’,
              E: ‘this.SELECT([I,D,E],\n’+
              ‘[this.gamma_E*(1-EXP(-this.gamma_E –
this.gamma_D))/(this.gamma_E+this.gamma_D),\n’+
              ‘this.gamma_D*(1-EXP(-this.gamma_E –
this.gamma_D))/(this.gamma_E+this.gamma_D),\n’+
              ‘EXP(-this.gamma_E - this.gamma_D)]);\n’,
              I: ‘this.SELECT([V,D,I],\n’+
              ‘[this.gamma_I*(1-EXP(-this.gamma_I –
this.gamma_ID))/(this.gamma_I+this.gamma_ID),\n’+
              ‘this.gamma_ID*(1-EXP(-this.gamma_I –
this.gamma_ID))/(this.gamma_I+this.gamma_ID),\n’+
              ‘EXP(-this.gamma_I - this.gamma_ID)]);\n’,
              V: ‘this.SELECT([S,D,V],\n’+
              ‘[this.gamma_V*(1-EXP(-this.gamma_V –
this.gamma_D))/(this.gamma_V+this.gamma_D),\n’+
              ‘this.gamma_D*(1-EXP(-this.gamma_V –
this.gamma_D))/(this.gamma_V+this.gamma_D),\n’+
              ‘EXP(-this.gamma_V - this.gamma_D)]);\n’,
              D: ‘D’,
              },
              },
              disease_trans : {
                 specifies: “InPin”,
                 exp: “mscope.disease_trans”,
              },
              disease_death : {
                 specifies: “InPin”,
                 exp: “mscope.disease_death”,
              },
              p_switch : {
                 specifies: “InPin”,
                 exp: “mscope.p_switch”,
              },
              nat_death : {
                 specifies: “InPin”,
                 exp: “mscope.nat_death”,
              },
              weight_res : {
                 specifies: “InPin”,
                 exp: “mscope.weight_res”,
              },
              weight_occ : {
                 specifies: “InPin”,
                 exp: “mscope.weight_occ”,
              },
              res_grow : {
                 specifies: “InPin”,
                 exp: “mscope.res_grow”,
              },
              comp_ex : {
                 specifies: “InPin”,
                 exp: “mscope.comp_ex”,
              },
              rep_fit_thresh : {
              pecifies: “InPin”,
                 exp: “mscope.rep_fit_thresh”,
              },
              res_def : {
                 specifies: “InPin”,
                 exp: “mscope.res_def”,
              },
              stress_decay : {
                 specifies: “InPin”,
                 exp: “mscope.stress_decay”,
              },
              stress_death : {
                 specifies: “InPin”,
                 exp: “mscope.stress_death”,
              },
              fit_decay : {
                 specifies: “InPin”,
                 exp: “mscope.fit_decay”,
              },
              move_cost : {
                 specifies: “InPin”,
                 exp: “mscope.move_cost”,
              },
           },
       });
  defineSchema(“main”, {
                 specifies: “Capsule”,
              methods: {
              },
              components: {
              NWorld : {
                 specifies: “SimWorld”,
                 count: 25,
                 rows: 18,
                 cols: 18,
                 agentbase: “Agent”,
                 cellbase: “Cell”,
                 motion: “discrete”,
                 layout: “hexagonal”,
                 ainputs: {
                    Init_Count: “this.Super.init_agent_count”,
                    CellNests: “this.Super.Nests”,
              },
              cinputs: {
                   Nests: “this.Super.Nests”,
              },
          },
              Nests : {
                   specifies: “Term”,
                   constant: true,
                   exp: ‘this.ASVALUE(\n’+
              ‘[“C3:3”,”C3:9”,”C3:15”,\n’+
              ‘“C9:3”,”C9:9”,”C9:15”,\n’+
              ‘“C15:3”,”C15:9”,”C15:15”\n’+
              ‘]\n’+
              ‘);\n’,
            },
            Agent_count : {
                 specifies: “Term”,
                 exp: ‘this.NWorld.AData(“AGENTCOUNT”)’,
            },
              colony_0 : {
                 specifies: “Term”,
                 exp: ‘nestPop[0]’,
            },
              colony_1 : {
                 specifies: “Term”,
                 exp: ‘nestPop[1]’,
            },
              colony_2 : {
                 specifies: “Term”,
                 exp: ‘nestPop[2]’,
            },
              colony_3 : {
                 specifies: “Term”,
                 exp: ‘nestPop[3]’,
            },
              colony_4 : {
                 specifies: “Term”,
                 exp: ‘nestPop[4]’,
            },
              colony_5 : {
                 specifies: “Term”,
                 exp: ‘nestPop[5]’,
            },
              colony_6 : {
                 specifies: “Term”,
                 exp: ‘nestPop[6]’,
            },
              colony_7 : {
                 specifies: “Term”,
                 exp: ‘nestPop[7]’,
            },
              colony_8 : {
                 specifies: “Term”,
                 exp: ‘nestPop[8]’,
           },
              initial_nest_count : {
                 specifies: “Command”,
                 when: “pre”,
                 exp: function(){
                    nests = this.Nests();
                    for (var i = 0; i < nests.length; i++) {
                      nestPop[i] =
this.NWorld.CELL(nests[i]).MYAGENT_COUNT();
              }
              }
          },
            Nest_Population : {
              specifies: “Plugin”,
              base: “PL_Barchart”,
              pins: {
                inpt0: “this.colony_0”,
                inpt1: “this.colony_1”,
                inpt2: “this.colony_2”,
                inpt3: “this.colony_3”,
                inpt4: “this.colony_4”,
                inpt5: “this.colony_5”,
                inpt6: “this.colony_6”,
                inpt7: “this.colony_7”,
                inpt8: “this.colony_8”,
              },
              properties: props [“main”][“Nest_Population”],
            },
            init_agent_count : {
                 specifies: “Slider”,
              properties: props [“main”][“init_agent_count”],
            },
            Stats_Stress : {
                 specifies: “Plugin”,
                 base: “PL_Stats”,
                 pins: {
                    source: “NWorld_vector”,
                    In: “stress_out”,
              },
              properties: props [“main”][“Stats_Stress”],
            },
            Stats_Stress_graph : {
                 specifies: “Plugin”,
                 base: “PL_Linechart”,
                 pins: {
                   inpt0: “this.Stats_Stress.Mean”,
                   inpt1: “this.Stats_Stress.M_Plus_S”,
                   inpt2: “this.Stats_Stress.M_Minus_S”,
              },
              properties: props [“main”][“Stats_Stress_graph”],
           },
           Stats_Fitness : {
                 specifies: “Plugin”,
                 base: “PL_Stats”,
                 pins: {
                  source: “NWorld_vector”,
                  In: “fitness_out”,
              },
              properties: props [“main”][“Stats_Fitness”],
           },
           Stats_Fitness_graph : {
                 specifies: “Plugin”,
                 base: “PL_Linechart”,
                 pins: {
                  inpt0: “this.Stats_Fitness.Mean”,
                  inpt1: “this.Stats_Fitness.M_Plus_S”,
                  inpt2: “this.Stats_Fitness.M_Minus_S”,
              },
              properties: props [“main”][“Stats_Fitness_graph”],
           },
              Disease_Tab : {
                 specifies: “Plugin”,
                 base: “PL_Tabulator”,
                 pins: {
                    source: “NWorld_vector”,
                  In: “Disease_Out”,
              },
              properties: props [“main”][“Disease_Tab”],
          },
              Disease_Tab_graph : {
                 specifies: “Plugin”,
                  base: “PL_Linechart”,
                 pins: {
                  inpt0: “this.Disease_Tab.Out_00”,
                  inpt1: “this.Disease_Tab.Out_01”,
                  inpt2: “this.Disease_Tab.Out_02”,
                  inpt3: “this.Disease_Tab.Out_03”,
                  inpt4: “this.Disease_Tab.Out_04”,
              },
              properties: props [“main”][“Disease_Tab_graph”],
           },
           Stats_Resource : {
                 specifies: “Plugin”,
                 base: “PL_Stats”,
                 pins: {
              source: “NWorld_matrix”,
              In: “resource_out”,
              },
              properties: props [“main”][“Stats_Resource”],
              },
              Stats_Resource_graph : {
                 specifies: “Plugin”,
                 base: “PL_Linechart”,
                 pins: {
                  inpt0: “this.Stats_Resource.Mean”,
                  inpt1: “this.Stats_Resource.M_Plus_S”,
                  inpt2: “this.Stats_Resource.M_Minus_S”,
              },
              properties: props [“main”][“Stats_Resource_graph”],
           },
           NWorld_display : {
                 specifies: “Plugin”,
                 base: “PL_AgentViewerX”,
                 pins: {
                  agentsource: “NWorld”,
                  cellsource: “NWorld”,
                  aColorOut: “agentColor”,
                  cColorOut: “cellColor”,
                  aSizeOut: “fitness”,
              },
              properties: props [“main”][“NWorld_display”],
           },
           Agent_count_spy : {
                 specifies: “Plugin”,
                 base: “PL_Spy”,
                 pins: {
                  inpt: “this.Agent_count”,
              },
              properties: props [“main”][“Agent_count_spy”],
           },
           disease_trans : {
                 specifies: “Slider”,
                 properties: props [“disease_trans”],
           },
           disease_death : { specifies: “Slider”,
                 properties: props [“disease_death”],
            },
            p_switch : {
                 specifies: “Slider”,
                 properties: props [“p_switch”],
            },
            nat_death : {
                 specifies: “Slider”,
                 properties: props [“nat_death”],
            },
           weight_res : {
                 specifies: “Slider”,
                 properties: props [“weight_res”],
           },
           weight_occ : { specifies: “Slider”,
                 properties: props [“weight_occ”],
           },
           res_grow : {
                 specifies: “Slider”,
                 properties: props [“res_grow”],
            },
           comp_ex : {
                 specifies: “Slider”,
                 properties: props [“comp_ex”],
            },
            rep_fit_thresh : {
                 specifies: “Slider”,
                 properties: props [“rep_fit_thresh”],
            },
            res_def : {
                 specifies: “Slider”,
                 properties: props [“res_def”],
            },
            stress_decay : {
                 specifies: “Slider”,
                 properties: props [“stress_decay”],
            },
            stress_death : {
                 specifies: “Slider”,
                 properties: props [“stress_death”],
            },
            fit_decay : {
                 specifies: “Slider”,
                 properties: props [“fit_decay”],
            },
            move_cost : {
                 specifies: “Slider”,
                 properties: props [“move_cost”],
            },
           },
        });
MAKE_ENUM(“Disease_State”,”S”,”E”,”I”,”V”,”D”);
MAKE_ENUM(“Action”,”stayed”,”moved”,”reproduced”);
MAKE_ENUM(“Sex”,”Male”,”Female”);
MAKE_ENUM(“Celltype”,”Nest”,”Normal”); 
var nestPop = [0,0,0,0,0,0,0,0,0]
             
var nestHSLs = [
              [30,100,50],
              [70,100,50],
              [110,100,50],
              [150,100,50],
              [190,100,50],
              [230,100,50],
              [270,100,50],
              [310,100,50],
              [350,100,50],
           ];
var clockParams = {
            lo: 0.000000, hi: 2000.000000, max: 2000.000000, dt: 1.000000,
          method: “Discrete”, step: 1, unit:”null”
              }
var props = {
            title: “sirTerrM”,
            address: “rms@cs.oberlin.edu”,
            date: “Sun Sep 08 08:49:02 PDT 2019”,
            paramset: “Param_set_1”,
            clockParams: {lo: 0.000000, hi: 2000.000000, max: 2000.000000, dt:
 1.000000, method: “Discrete”, step: 1, unit:”null”}, disease_trans: {
              type: “slider”, id: “disease_trans”, text: “disease_trans”, size: 1.000000,
              min: 0.000000, max: 1.000000, value: 0, step:
              0.010000, unit: ““
            },
              disease_death: {
              type: “slider”, id: “disease_death”, text: “disease_death”,
              size: 1.000000, min: 0.000000, max: 1.000000, value: 0, step:
              0.010000, unit: ““
            },
              p_switch: {
              type: “slider”, id: “p_switch”, text: “p_switch”, size:
              1.000000, min: 0.000000, max: 1.000000, value: 0, step: 0.100000,
              unit: ““
            },
              nat_death: {
              type: “slider”, id: “nat_death”, text: “nat_death”, size:
              1.000000, min: 0.000000, max: 0.100000, value: 0.005, step: 0.001000,
              unit: ““
            },
              weight_res: {
              type: “slider”, id: “weight_res”, text: “weight_res”, size:
              1.000000, min: 0.000000, max: 1.000000, value: 1/3, step: 0.010000,
              unit: ““
            },
              weight_occ: {
              type: “slider”, id: “weight_occ”, text: “weight_occ”, size:
              1.000000, min: 0.000000, max: 1.000000, value: 1/3, step: 0.010000,
              unit: ““
            },
              res_grow: {
              type: “slider”, id: “res_grow”, text: “res_grow”, size:
              1.000000, min: 0.000000, max: 1.000000, value: 0.2, step: 0.010000,
              unit: ““
            },
              comp_ex: {
              type: “slider”, id: “comp_ex”, text: “comp_ex”, size:
              1.000000, min: 0.000000, max: 20.000000, value: 1, step: 1.000000,
              unit: ““
            },
              rep_fit_thresh: {
              type: “slider”, id: “rep_fit_thresh”, text: “rep_fit_thresh”,
              size: 1.000000, min: 0.000000, max: 1.000000, value: 0.15, step:
              0.050000, unit: ““
            },
              res_def: {
              type: “slider”, id: “res_def”, text: “res_def”, size:
              1.000000, min: 0.000000, max: 0.500000, value: 0.1, step: 0.010000,
              unit: ““
            },
              stress_decay: {
              type: “slider”, id: “stress_decay”, text: “stress_decay”,
              size: 1.000000, min: 0.000000, max: 1.000000, value: 0.05, step:
              0.010000, unit: ““
            },
              stress_death: {
              type: “slider”, id: “stress_death”, text: “stress_death”, size: 1.000000, min:
              0.000000, max: 10.000000, value: 1, step:
              1.000000, unit: ““
            },
              fit_decay: {
              type: “slider”, id: “fit_decay”, text: “fit_decay”, size:
              1.000000, min: 0.000000, max: 1.000000, value: 0.1, step: 0.010000,
              unit: ““
            },
              move_cost: {
              type: “slider”, id: “move_cost”, text: “move_cost”, size:
              1.000000, min: 1.000000, max: 5.000000, value: 2, step: 0.200000,
              unit: ““
            },
              main: {
                  init_agent_count: {
                     pinned: false,
                     type: “slider”, id: “main:init_agent_count”, text: “agent count”, size: 6.000000, min: 0.000000, max: 1000.000000, value:
              350.000000, step: 50.000000, unit: ““
            },
            Nest_Population: {
              pinned: false,
              width: 500,
              height: 350,
              autogen: true,
              plotInterval: 1,
              contplot: true,
              XAxis: [“colony_0”, “colony_1”, “colony_2”, “colony_3”, “colony_4”, “colony_5”, “colony_6”, “colony_7”, “colony_8”],
              YAxis: [0.000000, 200.000000],
              caption: “Nest_Population”,
              strokes: [“#ff5500”, “#d4ff00”, “#2aff00”, “#00ff7f”,
              “#00d4ff”, “#002aff”, “#7f00ff”, “#ff00d4”, “#ff0026”, “#0000ff”,
              “#ff0000”, “#00ff00”, “#aaaa00”, “#ff00ff”, “#00ffff”, “#77777777”],
            },
              Stats_Stress: {
              input: “NWorld_vector.stress_out”,
            },
              Stats_Stress_graph: {
              pinned: false,
              width: 500,
              height: 350,
              autogen: true,
              multi: false,
              rate: 1,
              lo: clockParams .lo,
              hi: clockParams .hi,
              contplot: true,
              plotInterval: 10,
              tolerance: 0.001000,
              multipin: false,
              caption: “Stress Stats”,
              strokes: [“#00ff00”, “#0000ff”, “#ff0000”, “#aaaa00”, “#ff00ff”, “#00ffff”, “#77777777”],
              legend: [“Mean”, “Mean+Std”, “Mean-Std”],
              ylow: [null, null, null],
              yhigh: [null, null, null],
       },
       Stats_Fitness: {
              input: “NWorld_vector.fitness_out”,
       },
        Stats_Fitness_graph: {
              pinned: false,
              width: 500,
              height: 350,
              autogen: true,
              multi: false,
              rate: 1,
              lo: clockParams .lo,
              hi: clockParams .hi,
              contplot: true,
              plotInterval: 10,
              tolerance: 0.001000,
              multipin: false,
              caption: “Fitness Stats”,
              strokes: [“#00ff00”, “#0000ff”, “#ff0000”, “#aaaa00”, “#ff00ff”, “#00ffff”, “#77777777”],
              legend: [“Mean”, “Mean+Std”, “Mean-Std”],
              ylow: [null, null, null],
              yhigh: [null, null, null],
            },
             Disease_Tab: {
              input: “NWorld_vector.Disease_Out”,
              size: 5,
                 base: 0,
            },
             Disease_Tab_graph: {
              pinned: false,
              width: 500,
              height: 350,
              autogen: true,
              multi: false,
              rate: 1,
              lo: clockParams .lo,
              hi: clockParams .hi,
              contplot: true,
              plotInterval: 10,
              tolerance: 0.001000,
              multipin: false,
              caption: “Disease State”,
              strokes: [“#00ff00”, “#804d80”, “#e64d4d”, “#ff9966”, “#1a1a1a”, “#0000ff”, “#ff0000”, “#aaaa00”, “#ff00ff”, “#00ffff”, “#77777777”],
              legend: [“S”, “E”, “I”, “V”, “D”],
              ylow: [null, null, null, null, null],
              yhigh: [null, null, null, null, null],
            },
            Stats_Resource: {
              input: “NWorld_matrix.resource_out”,
            },
            Stats_Resource_graph: {
              pinned: false, width: 500,
              height: 350,
              autogen: true,
              multi: false,
              rate: 1,
              lo: clockParams .lo,
              hi: clockParams .hi,
              contplot: true,
              plotInterval: 10,
              tolerance: 0.001000,
              multipin: false,
              caption: “Resource Stats”,
              strokes: [“#00ff00”, “#0000ff”, “#ff0000”, “#aaaa00”, “#ff00ff”, “#00ffff”, “#77777777”],
              legend: [“Mean”, “Mean+Std”, “Mean-Std”],
              ylow: [null, null, null],
              yhigh: [null, null, null],
          },
           NWorld_display: {
              caption: “NWorld”,
              width: 540,
              height: 540,
              rows: 18,
              cols: 18,
              unit: 30,
              radius: 5,
              useAgent: true,
              useCell: true,
              interactive: false,
              agentSizing: true,
              largeSize: 5,
              smallSize: -1, colors: {
              cell: [“#000000”, “#ff0000”, “#008000”, “#0000ff”,
              “#00ffff”, “#ff00ff”, “#ffff00”, “#ffffff”],
              agent: [“#000000”, “#ff0000”, “#008000”, “#0000ff”,
              “#00ffff”, “#ff00ff”, “#ffff00”, “#ffffff”],
            },
              cellValues: [0.000, 1.000, 2.000, 3.000, 4.000, 5.000,
              6.000, 7.000],
              numAgentColors: 8,
              numCellColors: 8,
              staticCell: false,
              agentColorAssigned: false,
              hiAgentColor: 7.000000,
              loAgentColor: 0.000000,
              hiCellColor: 7.000000,
              loCellColor: 0.000000,
            },
              Agent_count_spy: {
              caption: “Agent_count”,
            },
         },
         Agent: {
         },
         Cell: {
          },
 };
~~~

## References

1. Dougherty ER, Seidel DP, Carlson CJ, Spiegel O, Getz WM. Going through the motions: incorporating movement analyses into disease research. Ecology letters. 2018;21(4):588–604.

2. Altizer S, Bartel R, Han BA. Animal migration and infectious disease risk. science. 2011;331(6015):296–302.

3. Altizer S, Ostfeld RS, Johnson PT, Kutz S, Harvell CD. Climate change and infectious diseases: from evidence to a predictive framework. science. 2013;341(6145):514–519.

4. Johnson PT, De Roode JC, Fenton A. Why infectious disease research needs community ecology. Science. 2015;349(6252):1259504.

5. Plowright RK, Sokolow SH, Gorman ME, Daszak P, Foley JE. Causal inference in disease ecology: investigating ecological drivers of disease emergence. Frontiers in Ecology and the Environment. 2008;6(8):420–429.

6. Smith KF, Dobson AP, McKenzie FE, Real LA, Smith DL, Wilson ML. Ecological theory to enhance infectious disease control and public health policy. Frontiers in Ecology and the Environment. 2005;3(1):29–37.

7. A’Bear AD, Jones TH, Boddy L. Size matters: what have we learnt from microcosm studies of decomposer fungus–invertebrate interactions? Soil Biology and Biochemistry. 2014;78:274–283.

8. Stewart RI, Dossena M, Bohan DA, Jeppesen E, Kordas RL, Ledger ME, et al. Mesocosm experiments as a tool for ecological climate-change research. In: Advances in ecological research. vol. 48. Elsevier; 2013. p. 71–181.

9. White LA, Forester JD, Craft ME. Disease outbreak thresholds emerge from interactions between movement behavior, landscape structure, and epidemiology. Proceedings of the National Academy of Sciences. 2018;115(28):7374–7379.

10. Hanski I, Gilpin ME, McCauley DE. Metapopulation biology. vol. 454. Elsevier; 1997.

11. Getz WM, Marshall CR, Carlson CJ, Giuggioli L, Ryan SJ, Romañach SS, et al. Making ecological models adequate. Ecology letters. 2018;21(2):153–166.

12. Larsen LG, Eppinga MB, Passalacqua P, Getz WM, Rose KA, Liang M. Appropriate complexity landscape modeling. Earth-science reviews. 2016;160:111–130.

13. Hethcote HW. The mathematics of infectious diseases. SIAM Review. 2000;42(4):599–653.

14. Anderson RM, May RM. Infectious diseases of humans: dynamics and control. Oxford university press; 1991.

15. Getz WM, Dougherty ER. Discrete stochastic analogs of Erlang epidemic models. Journal of biological dynamics. 2018;12(1):16–38.

16. Getz WM, Lloyd-Smith JO. Basic methods for modeling the invasion and spread of contagious diseases. DIMACS Series in Discrete Mathematics and Theoretical Computer Science. 2006;71:87.

17. Lloyd AL, Jansen VA. Spatiotemporal dynamics of epidemics: synchrony in metapopulation models. Mathematical biosciences. 2004;188(1):1–16.

18. Yang J, Liang S, Zhang Y. Travelling waves of a delayed SIR epidemic model with nonlinear incidence rate and spatial diffusion. Plos One. 2011;6(6):e21128.

19. Casagrandi R, Bolzoni L, Levin SA, Andreasen V. The SIRC model and influenza A. Mathematical biosciences. 2006;200(2):152–169.

20. Dushoff J, Plotkin JB, Levin SA, Earn DJ. Dynamical resonance can account for seasonality of influenza epidemics. Proceedings of the National Academy of Sciences. 2004;101(48):16915–16916.

21. Grenfell B, Bjørnstad O, Kappey J. Travelling waves and spatial hierarchies in measles epidemics. Nature. 2001;414(6865):716–723.

22. Lloyd-Smith JO, Galvani AP, Getz WM. Curtailing transmission of severe acute respiratory syndrome within a community and its hospital. Proceedings of the Royal Society of London B: Biological Sciences. 2003;270(1528):1979–1989.

23. Bauch CT, Lloyd-Smith JO, Coffee MP, Galvani AP. Dynamically modeling SARS and other newly emerging respiratory illnesses: past, present, and future. Epidemiology. 2005;16(6):791–801.

24. Althaus CL. Estimating the reproduction number of Ebola virus (EBOV) during the 2014 outbreak in West Africa. PLoS currents. 2014;6.

25. Getz WM, Gonzalez JP, Salter R, Bangura J, Carlson C, Coomber M, et al. Tactics and strategies for managing Ebola outbreaks and the salience of immunization. Computational and mathematical methods in medicine. 2015;2015.

26. Getz WM, Salter R, Muellerklein O, Yoon HS, Tallam K. Modeling epidemics: A primer and Numerus Model Builder implementation. Epidemics. 2018;.

27. Getz WM, Salter R, Mgbara W. Adequacy of SEIR models when epidemics have spatial structure: Ebola in Sierra Leone. Philosophical Transactions of the Royal Society B. 2019;374(1775):20180282.

28. Rivers CM, Lofgren ET, Marathe M, Eubank S, Lewis BL. Modeling the impact of interventions on an epidemic of Ebola in Sierra Leone and Liberia. PLoS currents. 2014;6.

29. Keeling MJ, Rohani P. Modeling infectious diseases in humans and animals. Princeton University Press; 2011.

30. Grimm V, Berger U, Bastiansen F, Eliassen S, Ginot V, Giske J, et al. A standard protocol for describing individual-based and agent-based models. Ecological modelling. 2006;198(1-2):115–126.

31. Lloyd-Smith JO, Schreiber SJ, Kopp PE, Getz WM. Superspreading and the effect of individual variation on disease emergence. Nature. 2005;438(7066):355.

32. Sah P, Leu ST, Cross PC, Hudson PJ, Bansal S. Unraveling the disease consequences and mechanisms of modular structure in animal social networks. Proceedings of the National Academy of Sciences. 2017;114(16):4165–4170. doi:10.1073/pnas.1613616114.

33. James AD, Rushton J. The economics of foot and mouth disease. Revue scientifique et technique-office international des epizooties. 2002;21(3):637–641.

34. Alexander KA, Carlson CJ, Lewis BL, Getz WM, Marathe MV, Eubank SG, et al. The Ecology of Pathogen Spillover and Disease Emergence at the Human-Wildlife-Environment Interface. In: The Connections Between Ecology and Infectious Disease. Springer; 2018. p. 267–298.

35. Carlson CJ, Getz WM, Kausrud KL, Cizauskas CA, Blackburn JK, Bustos Carrillo FA, et al. Spores and soil from six sides: interdisciplinarity and the environmental biology of anthrax (Bacillus anthracis). Biological Reviews. 2018;93(4):1813–1831.

36. De Garine-Wichatitsky M, Caron A, Kock R, Tschopp R, Munyeme M, Hofmeyr M, et al. A review of bovine tuberculosis at the wildlife–livestock–human interface in sub-Saharan Africa. Epidemiology & Infection. 2013;141(7):1342–1356.

37. Dean AS, Crump L, Greter H, Schelling E, Zinsstag J. Global burden of human brucellosis: a systematic review of disease frequency. PLoS neglected tropical diseases. 2012;6(10):e1865.

38. Williams E. Chronic wasting disease. Veterinary pathology. 2005;42(5):530–549.

39. Saadatnia G, Golkar M. A review on human toxoplasmosis. Scandinavian journal of infectious diseases. 2012;44(11):805–814.

40. Grimm V, Berger U, DeAngelis DL, Polhill JG, Giske J, Railsback SF. The ODD protocol: a review and first update. Ecological modelling. 2010;221(23):2760–2768.

41. Müller B, Bohn F, Dreßler G, Groeneveld J, Klassert C, Martin R, et al. Describing human decisions in agent-based models–ODD+ D, an extension of the ODD protocol. Environmental Modelling & Software. 2013;48:37–48.

42. Gotelli NJ, Graves GR. Null models in ecology. Smithsonian Institution; 1996.

43. Leibold MA. The niche concept revisited: mechanistic models and community context. Ecology. 1995;76(5):1371–1382.

44. Levin SA. Community equilibria and stability, and an extension of the competitive exclusion principle. The American Naturalist. 1970;104(939):413–423.

45. Armstrong RA, McGehee R. Competitive exclusion. The American Naturalist. 1980;115(2):151–170.

46. May RM, Mac Arthur RH. Niche overlap as a function of environmental variability. Proceedings of the National Academy of Sciences. 1972;69(5):1109–1113.

47. Pianka ER. Niche overlap and diffuse competition. Proceedings of the National Academy of Sciences. 1974;71(5):2141–2145.

48. Lloyd-Smith JO, Cross PC, Briggs CJ, Daugherty M, Getz WM, Latto J, et al. Should we expect population thresholds for wildlife disease? Trends in ecology & evolution. 2005;20(9):511–519.

49. Getz WM, Pickering J. Epidemic models: thresholds and population regulation. The American Naturalist. 1983;121(6):892–898.

50. Edwards AM, Phillips RA, Watkins NW, Freeman MP, Murphy EJ, Afanasyev V, et al. Revisiting Lévy flight search patterns of wandering albatrosses, bumblebees and deer. Nature. 2007;449(7165):1044.

51. Galvani AP. Epidemiology meets evolutionary ecology. Trends in Ecology & Evolution. 2003;18(3):132–139.

52. Grassly NC, Fraser C. Mathematical models of infectious disease transmission. Nature Reviews Microbiology. 2008;6(6):477.

53. Ostfeld RS, Glass GE, Keesing F. Spatial epidemiology: an emerging (or re-emerging) discipline. Trends in ecology & evolution. 2005;20(6):328–336.

54. Keesing F, Holt RD, Ostfeld RS. Effects of species diversity on disease risk. Ecology letters. 2006;9(4):485–498.

55. Archie EA, Luikart G, Ezenwa VO. Infecting epidemiology with genetics: a new frontier in disease ecology. Trends in Ecology & Evolution. 2009;24(1):21–30.

56. Hamby D. A review of techniques for parameter sensitivity analysis of environmental models. Environmental monitoring and assessment. 1994;32(2):135–154.

57. Christopher Frey H, Patil SR. Identification and review of sensitivity analysis methods. Risk analysis. 2002;22(3):553–578.

58. Borgonovo E, Plischke E. Sensitivity analysis: a review of recent advances. European Journal of Operational Research. 2016;248(3):869–887.

59. Pianosi F, Beven K, Freer J, Hall JW, Rougier J, Stephenson DB, et al. Sensitivity analysis of environmental models: A systematic review with practical workflow. Environmental Modelling & Software. 2016;79:214–232.

60. Caswell H. Matrix population models. Encyclopedia of Environmetrics. 2006;3.

61. Coulson T, Catchpole EA, Albon SD, Morgan BJ, Pemberton J, Clutton-Brock TH, et al. Age, sex, density, winter weather, and population crashes in Soay sheep. Science. 2001;292(5521):1528–1531.

62. Diekmann O, Heesterbeek JAP. Mathematical epidemiology of infectious diseases: model building, analysis and interpretation. vol. 5. John Wiley & Sons; 2000.

63. Getz WM. Biomass transformation webs provide a unified approach to consumer–resource modelling. Ecology letters. 2011;14(2):113–124.

64. Getz WM, Owen-Smith N. Consumer-resource dynamics: quantity, quality, and allocation. PloS one. 2011;6(1):e14539.

65. Beddington JR. Mutual interference between parasites or predators and its effect on searching efficiency. The Journal of Animal Ecology. 1975; p. 331–340.

66. DeAngelis DL, Goldstein R, O’neill R. A model for tropic interaction. Ecology. 1975;56(4):881–892.

67. Giuggioli L, Kenkre V. Consequences of animal interactions on their dynamics: emergence of home ranges and territoriality. Movement ecology. 2014;2(1):20.

68. Blackburn JK, Ganz HH, Ponciano JM, Turner WC, Ryan SJ, Kamath P, et al. Modeling R0 for Pathogens with Environmental Transmission: Animal Movements, Pathogen Populations, and Local Infectious Zones. International journal of environmental research and public health. 2019;16(6):954.

69. Getz WM, Salter R, Lyons AJ, Sippl-Swezey N. Panmictic and clonal evolution on a single patchy resource produces polymorphic foraging guilds. PloS One. 2015;10(8):e0133732.

70. Getz WM, Salter R, Seidel DP, Van Hooft P. Sympatric speciation in structureless environments. BMC evolutionary biology. 2016;16(1):50.

71. Huisman G, De Boer RJ. A formal derivation of the “Beddington” functional response. Journal of theoretical biology. 1997;185(3):389–400.

72. Cantrell RS, Cosner C. On the dynamics of predator–prey models with the Beddington–DeAngelis functional response. Journal of Mathematical Analysis and Applications. 2001;257(1):206–222.

73. Getz WM. Computational population biology: linking the inner and outer worlds of organisms. Israel journal of ecology & evolution. 2013;59(1):2–16.

74. Ball F, Nåsell I. The shape of the size distribution of an epidemic in a finite population. Mathematical biosciences. 1994;123(2):167–181.

75. Gordillo LF, Marion SA, Martin-Löf A, Greenwood PE. Bimodal epidemic size distributions for near-critical SIR with vaccination. Bulletin of Mathematical Biology. 2008;70(2):589–602.

76. Lloyd-Smith JO, George D, Pepin KM, Pitzer VE, Pulliam JR, Dobson AP, et al. Epidemic dynamics at the human-animal interface. Science. 2009;326(5958):1362–1367.

77. Méléard S, Villemonais D, et al. Quasi-stationary distributions and population processes. Probability Surveys. 2012;9:340–410.

78. Fan M, Kuang Y. Dynamics of a nonautonomous predator–prey system with the Beddington–DeAngelis functional response. Journal of Mathematical Analysis and Applications. 2004;295(1):15–39.

79. Hwang TW. Global analysis of the predator–prey system with Beddington–DeAngelis functional response. Journal of Mathematical Analysis and Applications. 2003;281(1):395–401.

80. Li H, Takeuchi Y. Dynamics of the density dependent predator–prey system with Beddington–DeAngelis functional response. Journal of Mathematical Analysis and Applications. 2011;374(2):644–654.

81. Chen W, Wang M. Qualitative analysis of predator-prey models with Beddington-DeAngelis functional response and diffusion. Mathematical and Computer Modelling. 2005;42(1-2):31–44.

82. Huang G, Ma W, Takeuchi Y. Global properties for virus dynamics model with Beddington–DeAngelis functional response. Applied Mathematics Letters. 2009;22(11):1690–1693.

83. Huang G, Ma W, Takeuchi Y. Global analysis for delay virus dynamics model with Beddington–DeAngelis functional response. Applied Mathematics Letters. 2011;24(7):1199–1203.

84. Elaiw A, Azoz S. Global properties of a class of HIV infection models with Beddington–DeAngelis functional response. Mathematical Methods in the Applied Sciences. 2013;36(4):383–394.

85. Wangersky PJ. Lotka-Volterra population models. Annual Review of Ecology and Systematics. 1978;9(1):189–218.

86. Thierry H, Sheeren D, Marilleau N, Corson N, Amalric M, Monteil C. From the Lotka–Volterra model to a spatialised population-driven individual-based model. Ecological modelling. 2015;306:287–293.

87. Karsai I, Montano E, Schmickl T. Bottom-up ecology: an agent-based model on the interactions between competition and predation. Letters in Biomathematics. 2016;3(1):161–180.

88. Lofgren E, Fefferman NH, Naumov YN, Gorski J, Naumova EN. Influenza seasonality: underlying causes and modeling theories. Journal of virology. 2007;81(11):5429–5436.

89. Ovaskainen O, Meerson B. Stochastic models of population extinction. Trends in ecology & evolution. 2010;25(11):643–652.

90. Getz W, Muellerklein O, Salter R, Carslon C, Lyons A, Seidel D. A web app for population viability and harvesting analyses. Natural Resource Modeling. 2017;30(2).

91. McLane AJ, Semeniuk C, McDermid GJ, Marceau DJ. The role of agent-based models in wildlife ecology and management. Ecological Modelling. 2011;222(8):1544–1556.

92. Filatova T, Verburg PH, Parker DC, Stannard CA. Spatial agent-based models for socio-ecological systems: challenges and prospects. Environmental modelling & software. 2013;45:1–7.

## References

[1] Getz WM, Salter R, Lyons AJ, Sippl-Swezey N. Panmictic and clonal evolution on a single patchy resource produces polymorphic foraging guilds. PloS One. 2015;10(8):e0133732.

[2] Getz WM, Salter R, Muellerklein O, Yoon HS, Tallam K. Modeling epidemics: A primer and Numerus Model Builder implementation. Epidemics. 2018;.

